# The effect of gene tree dependence on summary methods for species tree inference

**DOI:** 10.1101/2024.06.06.597697

**Authors:** Wanting He, Celine Scornavacca, Yao-ban Chan

## Abstract

When inferring the evolutionary history of species and the genes they contain, the phylogenetic trees of genes can be different from those of the species and to each other, due to a variety of causes, including incomplete lineage sorting. We often wish to infer the species tree, but only reconstruct the gene trees from sequences. We then combine the gene trees to produce a species tree; methods to do this are known as summary methods, of which ASTRAL is currently among the most popular. ASTRAL has been shown to be accurate in many practical scenarios through extensive simulations. However, these simulations generally assume that the input gene trees are independent of each other (infinite recombination between loci). This is known to be unrealistic, as genes that are close to each other on the chromosome (or are co-evolving) have dependent phylogenies.

In this paper, we develop a model for generating dependent gene trees within a species tree, based on the coalescent with recombination. We then use these trees as input to ASTRAL to reassess its accuracy for dependent gene trees. Our results allow us to evaluate the impact of any level of dependence on the accuracy of ASTRAL, both when gene trees are known and estimated from sequences. We find that a fixed amount of dependence reduces the effective sample size by a constant factor.

In current phylogenomic datasets, loci are generally sampled at large genomic distances to reduce gene tree dependence, thereby limiting the number of genes available for inference. However, full independence between genes is not required for accurate species tree estimation, and excluding gene trees may reduce inference accuracy. This creates a trade-off between the number of genes used and the degree of gene tree dependence. We therefore propose a method to identify the minimum genomic separation required to maintain satisfactory inference accuracy.

## Introduction

Speciation is an evolutionary process where populations evolve and become distinct species (Coyne and Orr, 2004). A species tree, or phylogeny (Hillis et al., 1996), depicts the history of speciation where leaves represent extant species, internal nodes represent speciation events, and evolutionary distances are represented by branch lengths. Likewise, gene trees depict the evolutionary history of gene families within the genomes of these species. Species trees and gene trees play a vital role in the study of gene and genome evolution, and their reconstruction can give us insight into the history of life on Earth (Darwin, 1859). When species lineages diverge through speciation, gene copies within these species also diverge. Therefore, gene trees can be thought of as evolving within the branches of the species tree. However, in addition to speciation, various gene-only evolutionary processes can cause gene trees to be distinct from the species tree; these include gene duplication and loss, horizontal gene transfer (HGT), and incomplete lineage sorting (ILS; Maddison, 1997). In particular, ILS, where multiple gene lineages do not coalesce over a series of speciations, is a major cause of discordance between gene and species trees (see Figure 1 for an example). The standard statistical model for gene trees under ILS is the multispecies coalescent (MSC) model (Pamilo and Nei, 1988; Rannala and Yang, 2003). In this model, each species branch represents a separate population in which Kingman’s basic coalescent (Kingman, 1982) is run in a bottom-up fashion, and lineages at the top of each branch are used as input to their parent branches.

**Figure 1:**
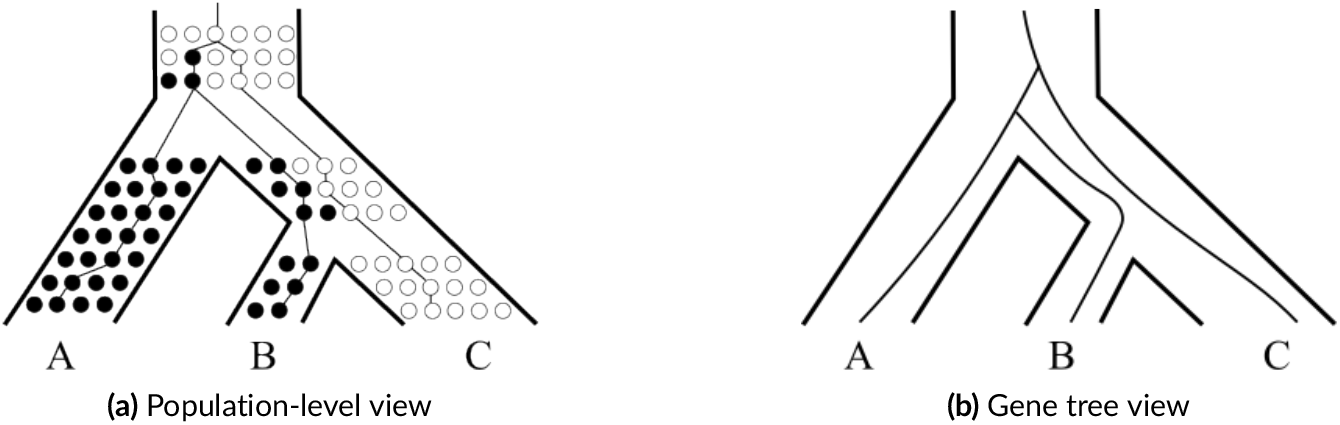
An example of incomplete lineage sorting. Circles represent individuals of a population, and each row corresponds to a generation. Originally, the population contains only individuals with an ancestral allele (open circles), and then a mutation introduces a derived allele (filled circles), creating genetic polymorphism. The ancestral allele does not descend in species *A*, leaving only individuals with the derived allele. The polymorphism persists in the ancestor of species *B* and *C*. Subsequently species *B* retains only the derived allele, while species *C* retains only the ancestral allele. For this gene family, individuals in population *B* are more closely related to individuals in population *A* than *C*, leading to topological discordance between the gene and species trees.

Because gene trees can be different from the species tree, we need to utilise multiple gene families to infer the species tree. A common method used in the past was to concatenate sequences from multiple genes to form a supermatrix, and then obtain a species tree by applying traditional phylogenetic inference methods, such as distance-based or maximum-likelihood. This paradigm implicitly assumes that all genes come from a single gene tree, which (for the reasons given above) is known not to be true.

More recently, summary methods have been developed, where sequences from different loci are analysed separately to build gene trees, which are then summarised in some way to generate a species tree. Some summary methods only use gene tree topologies, for example, MP-EST (Liu et al., 2010), NJst (Liu and Yu, 2011), ASTRID (Vachaspati and Warnow, 2015), DISTIQUE (Sayyari and Mirarab, 2016), ASTRAL (Mirarab et al., 2014b; Mirarab and Warnow, 2015; Zhang et al., 2018) and STAR (Liu et al., 2009), and some use both gene tree topologies and branch lengths, such as GLASS (Mossel and Roch, 2008) and STEAC (Liu et al., 2009). The input of these methods are mainly rooted gene trees, but some (such as ASTRAL, NJst, ASTRID, and DISTIQUE) can take unrooted gene trees as input. A number of other paradigms are also available, including full-likelihood (Minh et al., 2020; Nguyen et al., 2015; Stamatakis, 2014), Bayesian (Flouri et al., 2018; Heled and Drummond, 2009; Ogilvie et al., 2017; Rannala and Yang, 2017), and co-estimation methods (Heled and Drummond, 2009; Liu, 2008; Ogilvie et al., 2017).

ASTRAL is a popular summary method due to its high accuracy (Ballesteros and Sharma, 2019; Giarla and Esselstyn, 2015) and scalability (Mirarab and Warnow, 2015; Yin et al., 2019). It has been shown theoretically to give a statistically consistent estimator of the species tree if input gene trees are sampled under the MSC model (Mirarab et al., 2014b), bounded HGT models (Davidson et al., 2015), the general DLCoal model (Markin and Eulenstein, 2021), or the GDL model (Legried et al., 2021; Yan et al., 2022). Furthermore, extensive simulations have been performed studying its accuracy, showing that ASTRAL is highly accurate under the MSC model (Mirarab et al., 2014b; Mirarab and Warnow, 2015; Sayyari and Mirarab, 2016; Vachaspati and Warnow, 2015), the presence of both HGT and ILS (Davidson et al., 2015), and with gene filtering when ILS is low to moderate (Molloy and Warnow, 2018). An important practical consideration is whether the gene trees are known exactly or are estimated from sequences with some potential error. It is well known that the accuracy of all summary methods, including ASTRAL, is affected by gene tree estimation error (DeGiorgio and Degnan, 2014; Huang and Knowles, 2016; Lanier and Knowles, 2015; Molloy and Warnow, 2018; Patel et al., 2013).

Although the accuracy of ASTRAL has been extensively studied in simulations, they were almost all performed with gene trees that were simulated independently from the model of choice. In effect, this assumes no recombination within a gene and unlimited recombination between genes, both of which are known not to be realistic. The former assumption has been tested (Lanier and Knowles, 2012; Zhu et al., 2022) and found to have relatively little practical effect on the accuracy of species tree inference. However, these studies explored only a limited range of simulation scenarios, and more complex cases involving short internal branches, high substitution rate variation, and deep divergences among taxa remain largely untested.

In this paper, we focus on the latter assumption; in reality, gene trees are dependent because genes can be located near to each other on the same chromosome, which causes them to have related histories. Indeed, in the absence of recombination, adjacent genes will have identical phylogenies. Recombination breaks this dependence when and where it occurs, but because it occurs at a finite rate, the gene trees of nearby genes are not fully independent. Other causes, such as functional gene linkage, may also induce dependence between gene trees (Barker and Pagel, 2005). Thus, by sampling gene trees from an independent model, previous simulations may have mis-estimated the accuracy of ASTRAL.

This effect was previously studied by Wang and Liu (2016), who found a significant impact on the accuracy of ASTRAL when gene trees are dependent. However, their methodology attempted to delineate loci from whole genomes using inferred or known breakpoints; this has the effect of very strong dependence between neighbouring gene trees, which may not always occur. Additionally, Conry (2020) found that recombination between exons has little effect on the accuracy of species tree inference, but from simulations only on four-taxon species trees; the effect may become more noticeable with larger trees.

The effect of recombination on the evolutionary history of a genome is well-known in population genetics, where the standard statistical model for a single population is the coalescent with recombination (Hudson, 1983). This produces an evolutionary history that contains both coalescent and recombination events, known as the ancestral recombination graph (ARG; Griffiths and Marjoram, 1997). The program `ms` (Hudson, 2002) was an early tool for simulating the ARG; more recently, `msprime` (Baumdicker et al., 2022; Kelleher et al., 2016) was developed as a high-speed, large-scale successor, offering both rapid simulation and efficient data storage.

Phylogenomic data can be produced in many different ways. A common approach is to identify orthogolous gene families between the species of interest and extract the gene sequences. This results in full sequences that have some amount of genomic distance between them, as typically only exons are extracted. The dependence between the genes depends on the amount of separation; ideally the distance is sufficient to achieve approximately independent genes, but this is not always checked and the conditions for achieving a sufficient distance are not well studied. An alternative approach is to analyse the SNPs common to the sequences and separate them into loci in some way, often using linkage disequilibrium-based (LD) approaches to ensure approximately independent loci. As SNPs are evenly distributed among the genome, this approach discards some data to ensure sufficient separation.

Our aim in this paper is to reassess the accuracy of ASTRAL under conditions when gene trees are dependent. We generalise the two-locus, 3-taxon model of Slatkin and Pollack (2006), and derive a probabilistic method for generating dependent gene trees for multiple species and loci. We then use the generated trees as input to ASTRAL to estimate the species tree. This allows us to numerically estimate the impact of a number of factors, including the number of loci, the amount of ILS, recombination rate, and the dependence structure, on the accuracy of ASTRAL. For real data, we study approaches to ensuring that gene dependence does not affect the accuracy of species tree inference. We re-analyse an 8-taxon mouse SNP dataset (Liu et al., 2015), and find that satisfactory accuracy can be achieved by sampling SNPs at distances much shorter than those suggested by LD-based methods. For datasets produced by extracting orthologous genes, we devise a method to determine the minimum genomic distance in order to maintain satisfactory accuracy. The real-life 37-taxon mammalian dataset that we study (Mirarab et al., 2014b) has genes separated by much larger distances than this minimum, indicating that current datasets are minimally affected by gene tree dependence.

## Methods

### Generating dependent gene trees

To generate dependent gene trees within a species tree, we extend the model of Slatkin and Pollack (2006), which considered two loci of haploid individuals in a single Wright-Fisher (panmictic) population. In that paper, they devised a probabilistic model of two linked loci in three species based on the coalescent with recombination, using it to calculate the probabilities that the gene trees in the loci are concordant with the species tree and with each other. Here, we extend the model in a natural way to multiple species and loci, but more importantly use it to produce an algorithm to simulate a gene tree in one locus conditional on a known tree in a linked locus. This idea was also used in Li et al. (2021) to model so-called ‘linked duplications’. Figure 2 shows an example of our model (described below).

**Figure 2:**
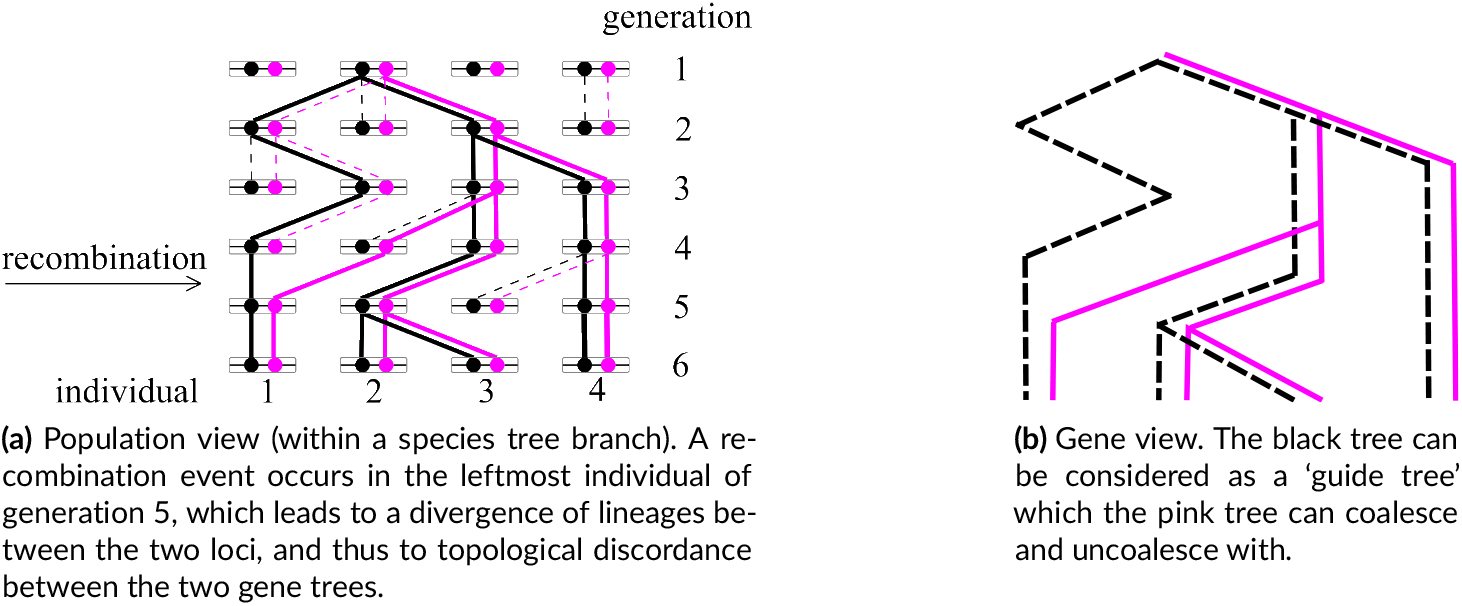
An example of the two-locus model. The black and pink lines represent the genealogies in the first and second loci respectively within a population (a species branch), and the black tree is considered as the guide tree.

Consider first a single panmictic population. In the absence of recombination, the parents of all individuals are the same between the loci, and so the genealogies will be the same. We model this by considering the known genealogy in the first locus as a ‘guide tree’, to which lineages in the second locus can ‘coalesce’, representing that the lineages belong to the same individual. If a lineage is coalesced with the guide tree, it must then follow the ancestry of that lineage if no recombination occurs. Because we observe extant samples of multiple loci in the same individuals, the lineages in the second locus begin coalesced with the guide tree at the present time. Thus, in the absence of recombination, the two trees will be identical.

Recombinations occur between the two loci at a rate of *R* per individual per coalescent unit. (If there is a genomic distance of *d* sites between loci, then in population genetics notation, *R* = 2*N*_*e*_ *rd*, where 2*N*_*e*_ is the effective population size and *r* is the recombination rate per site per generation.) When a recombination occurs, the two loci in an individual have distinct parents in the preceding generation. We represent this as the lineage in the second locus ‘uncoalescing’ from the guide tree, so that it no longer follows the ancestry of the first locus. However, recombination is an event that happens to a single individual/lineage, so other lineages are unaffected. Note in particular that coalesced lineages in the second locus cannot uncoalesce from each other, as this does not represent recombination.

Proceeding backwards in time, uncoalesced lineages in the second locus can either coalesce with a lineage of the guide tree (whether or not that lineage is already coalesced with a lineage in the second locus) or with each other. These coalescences occur at rates consistent with the multispecies coalescent. Thus if there are *k*_*u*_ uncoalesced lineages and *k*_*g*_ guide tree lineages in a population at a particular time, the uncoalesced lineages will coalesce with each other at a rate of 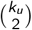 and with a guide tree lineage at a rate of *k*_*u*_*k*_*g*_ (in coalescent units). The times of coalescences between two guide tree lineages are specified by the guide tree. As with the basic coalescent, this process continues until all lineages in the second locus have reached their most recent common ancestor (MRCA), which is not necessarily the MRCA in the first locus. We extend this process to multiple species in a standard way.

This allows us to generate a gene tree conditional on another gene tree. To generate multiple dependent gene trees along a linear genome, we first generate a gene tree under the MSC. Each gene tree is then used as a guide tree in our model for the subsequent gene tree. For a linear genome, this reduces to the sequentially Markov coalescent (SMC; McVean and Cardin, 2005), sampled at constant intervals along the genome. However, our model is more flexible, as it can also simulate dependent gene trees where there is a non-linear dependence structure between genes (for example, when dependence is produced by functional rather than proximity relationships). As far as we know, our algorithm is the only one that can achieve this, although we do not use this functionality in this paper.

We note that there already exist several tools for simulating gene trees under the full coalescent with recombination, such as `ms` and `msprime`. We have elected not to use these tools, as `ms` is too slow (Chen et al., 2009), while `msprime` does not currently accept non-ultrametric species trees, which we have. Although our model lacks the long-range dependence structure of the full coalescent with recombination, it is faster to simulate, and as described above retains more flexibility. Nevertheless, we also conduct some simulations on ultrametric trees to determine that the results from our model do not differ significantly from those produced using gene trees generated by `msprime` (see Comparison with `msprime`).

The described model assumes a constant amount of recombination between loci, which is not biologically realistic due to varying gene positions and recombination rates along the genome. We also explore an alternative way of generating dependent gene trees where we generate independent ‘blocks’ of gene trees, where block boundaries are determined randomly along the genome. Within each block, we generate trees with a constant recombination rate *R* as above.

We perform this scenario in two different ways. First, we keep the recombination rate *R* fixed and vary the number of independent blocks, while keeping the total number of gene trees fixed. This also varies the ‘overall’ recombination rate, so to observe the effects for a fixed overall recombination rate, we use a second scheme in which we vary the recombination rate within each block while fixing the average value of 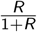 between trees. An intuitive explanation for this expression is that lineages coalesce at a rate of 1 and uncoalesce with a rate of *R*; therefore 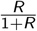 is approximately the fraction of time that a new tree spends uncoalesced from the guide tree, and hence quantifies the amount of new information each tree adds to the sample.

### Simulations

We explore the performance of ASTRAL using simulations on a 37-taxon mammalian species tree, which has previously been used to study the accuracy of ASTRAL for independent gene trees (Mirarab et al., 2014b). The species tree was previously estimated with MP-EST (Liu et al., 2010) on the biological dataset from Song et al. (2012), containing 447 genes with average length 3099bp. We re-estimate the branch lengths as specified in Supporting information, while keeping the same topology. The result is shown in Figure 3.

**Figure 3:**
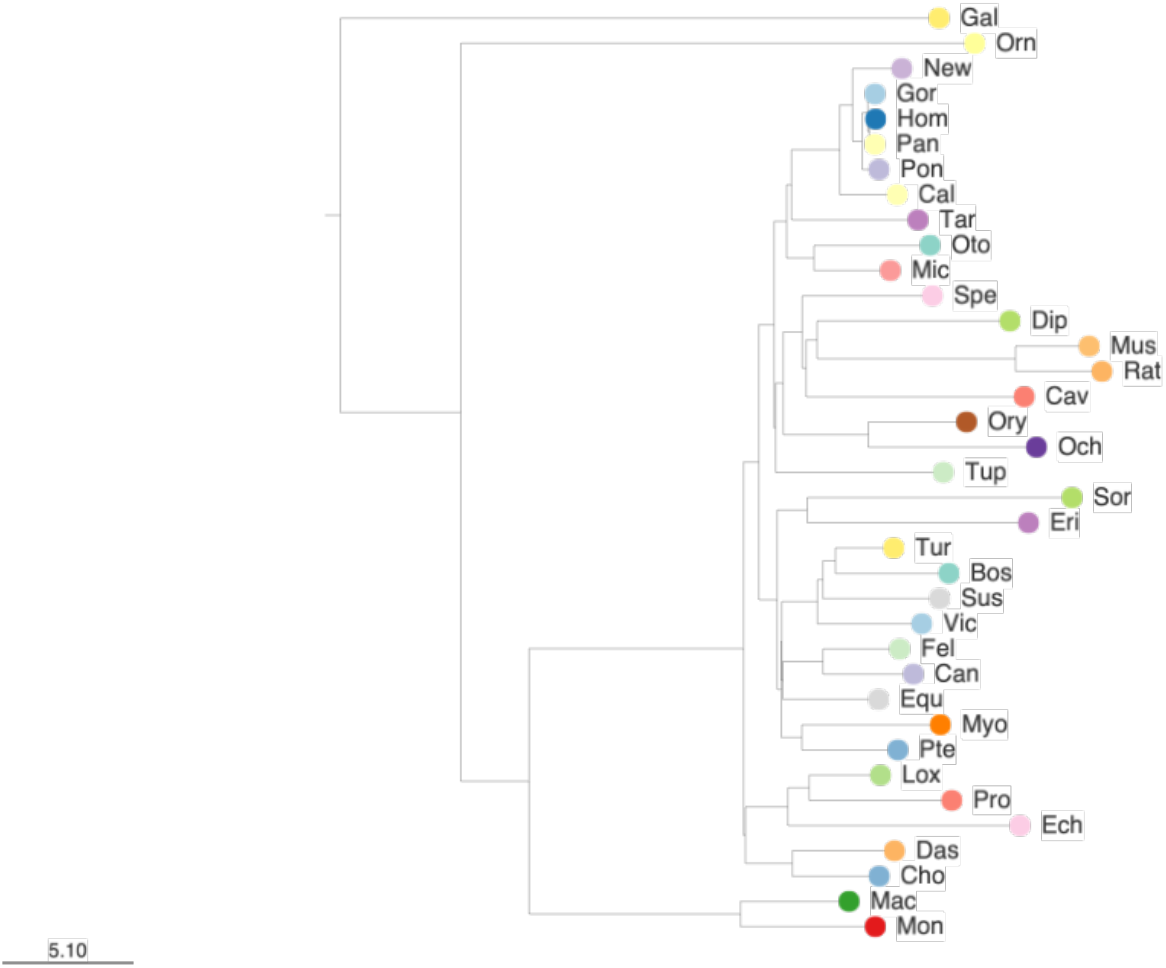
**The 37-taxon mammalian species tree (with branch lengths in coalescent units**) (full species names are given in Table 15 in Supporting information).

We perform simulations with the same parameter settings as in Mirarab et al. (2014b), adding dependence between the gene trees as specified above. We then use these trees as input to ASTRAL to infer a species tree, and evaluate the inference accuracy using the normalised Robinson-Foulds (RF) distance (Robinson and Foulds, 1981). The RF distance measures the difference between the true and inferred species trees by counting the number of clades present in one tree but not the other.

Additionally, we rescale the simulated dependent gene trees into units of substitutions per site, and then evolve gene sequences along the trees with `Pyvolve` (Spielman and Wilke, 2015), under the GTR+Г model of site evolution (Tavaré, 1986) with parameters estimated from the biological dataset (see Supporting information for more details). As in Mirarab et al. (2014b), we use three sequence lengths: 500bp, 1000bp, and a mixture where half of the sequences are 500bp and the other half are 1000bp. We then use IQ-TREE (Minh et al., 2020; Nguyen et al., 2015) to estimate gene trees from these sequences, and use the estimated gene trees as input to ASTRAL. This allows us to study the effect of gene tree estimation error.

From initial results, we found that accuracy varies the most when the recombination rate *R* lies between 0 and 1. Although this range is much smaller than the values observed in the dataset (see Section Estimating minimum genomic distance in a phylogenomic dataset for details), we nevertheless focus on it, as it captures the regime in which accuracy is most sensitive to changes in *R*. When *R* is equal to 0, all gene trees are identical; as it increases, there is less dependence between neighbouring gene trees, and when it is infinite, the gene trees are independent.

It has been shown that ASTRAL performs worse with an increased amount of ILS (Mirarab et al., 2014b), so we are also interested in the performance of ASTRAL with dependent gene trees in this case. We multiply the branch lengths of the species tree by 0.2 (denoted by 0.2×), which is equivalent to multiplying the effective population size by 5, and repeated the simulation above. Finally, we also fix the number of gene trees *N* to be 200, and evaluate the accuracy of ASTRAL with different amounts of ILS, with branch length multipliers ranging from 0.2× to 5× as in Mirarab et al. (2014b).

The parameter settings for the simulations are summarised in Table 1. We perform 100 replicates for each parameter setting for true gene trees, and 20 replicates for estimated gene trees (as gene tree estimation is computationally intensive). In addition, we only use *N* ≤ 800 gene trees per dataset for estimated gene trees. The same simulation settings are used when varying recombination rate along the genome, except that we use 100 replicates and fix the overall recombination rate (as described above) to be 0.1.

**Table 1:** Simulation parameters.

| Simulation | $R$ | $N$ | ILS<sup>1</sup> |
| --- | --- | --- | --- |
| Default simulations | [0, 1] | [100, 3200] | $1\times$ branch lengths |
| Increased amount of ILS | [0, 1] | [100, 3200] | $0.2\times$ branch lengths |
| Different amounts of ILS | [0, 1] | 200 | $[0.2, 5]\times$ branch lengths |

<sup>1</sup> Multiplying branch lengths of the species tree by $\alpha$ is equivalent to dividing the effective population size by $\alpha$ .

To verify that the our results are not specific to the species tree used, we repeat the above simulations (for true gene trees only) on a tree of 16 fungi (Butler et al., 2009; Legried et al., 2021; Markin and Eulenstein, 2021; Rasmussen and Kellis, 2012; Wu et al., 2014), shown in Figure 19 in Supporting information. This fungal tree is ultrametric and has been studied extensively in simulations aiming to show that the accuracy of ASTRAL is not affected by data with paralogs (Yan et al., 2022). For this dataset, we vary *R* from 0 to 0.5 following an initial exploration. Otherwise, we use the same simulation settings shown in Table 1.

To compare the performance of our model with `msprime` (Baumdicker et al., 2022; Kelleher et al., 2016), which is restricted to ultrametric species trees, we also conducted simulations under `msprime` on the fungal tree, using the same parameter settings, and compared these results to those generated from our model. Although the 37-taxon mammalian species tree we study is not ultrametric, we extend the terminal branches to obtain an ultrametric tree, and compare our model with `msprime`.

### Comparison with Wang and Liu

Wang and Liu (2016) also studied the effect of gene tree dependence on the accuracy of ASTRAL. To do so, they generated gene trees (and then sequences) under the (multispecies) coalescent with recombination with `ms` (Hudson, 2002). As they also studied the effect of breakpoint inference on ASTRAL, they compared five cases:

1. LD1000: Loci of 1000bp were sampled along the sequences at intervals estimated to be (near-)independent from linkage disequilibrium plots; gene trees were estimated from the loci with FastTree (Price et al., 2009, 2010).
2. LD100: As above, but with locus lengths of 100bp.
3. IBIG (inferred breakpoints, inferred gene trees): Recombination breakpoints were inferred and used to partition the sequences. Gene trees were then estimated from the partitions with FastTree.
4. TBIG (true breakpoints, inferred gene trees): As above, but known breakpoints (from `ms`) were used.
5. TBTG (true breakpoints, true gene trees): True gene trees (from `ms`) were used.

Apart from the model difference (which, as shown in Robustness of our results, is not significant), the two major differences between our study and the TBIG/TBTG cases of Wang and Liu (2016) are:

- They use the entire sequences, and partition at every breakpoint; thus their gene trees are separated by exactly one recombination (and therefore highly dependent).
- They do not explicitly control the number of gene trees, and do not directly compare with (the same number of) independent gene trees.

By varying the recombination rate between gene trees, we are able to study scenarios of lesser dependence (such as might occur when we recognise orthologous genes rather than aligning and partitioning whole-genome sequences). By controlling the number of gene trees, we are able to directly compare with independent trees, and thus quantify the effect of gene tree dependence.

## Results

### Default simulations

For the first simulation, we study the accuracy of ASTRAL in relation to the recombination rate between gene trees *R* and the number of input gene trees *N*. We show the results for both true and estimated gene trees in Figure 4.

**Figure 4:**
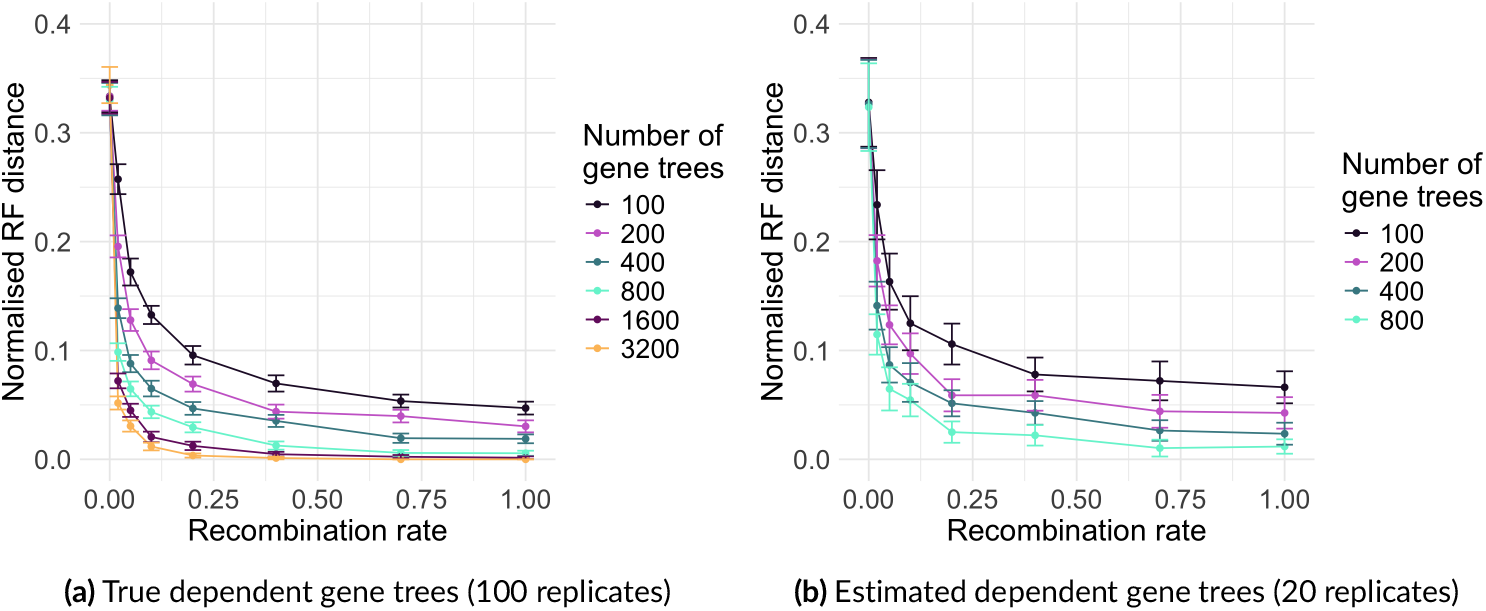
The accuracy of ASTRAL vs recombination rate, averaged over a number of replicates. Error bars represent standard error.

In Figure 4a, we show the results for true gene trees. When *R* = 0, all gene trees are identical, so the inferred species tree is equal to the single gene tree. Thus the accuracy of ASTRAL is unaffected by the number of input gene trees. As *R* increases, the normalised RF distance decreases — ASTRAL becomes more accurate with less dependence among gene trees. As *R* → ∞, the accuracy converges to the results of Mirarab et al. (2014b) for independent gene trees.

For constant *R*, ASTRAL becomes more accurate as the number of gene trees *N* increases, which is the same as for independent gene trees. It has been shown (Chan et al., 2022) that with independent gene trees, the normalised RF distance decreases exponentially with respect to *N*. In Figure 21 in Supporting information, we show a log-plot of normalised RF distance against *N*. It is inconclusive from this plot if the exponential asymptotic behaviour still holds for finite recombination rate; potentially it still does, but may take longer to reach the limiting behaviour.

Figure 4b shows the results for estimated gene trees from mixed sequence lengths, with detailed results for 500bp and 1000bp available in Figure 16 Supporting information. The three sequence lengths show minimal differences. In Figure 4b, we see that similar patterns hold when gene trees are estimated from sequences. Overall, the accuracy is slightly lower for estimated gene trees at all recombination rates, as expected. This indicates that the accuracy of ASTRAL can be adversely affected simultaneously by both gene tree estimation error and gene tree dependence. However, in the regime under study (*R <* 1), the effect of gene tree dependence is much larger than gene tree estimation error.

### Increased amount of ILS

We repeat our simulations above with an increased amount of ILS (0.2× branch lengths). We observe similar patterns as in the Default simulations section, as shown in Figure 5, with results for 500bp and 1000bp sequence lengths in Figure 17 Supporting information. As before, ASTRAL becomes more accurate with less dependence and with more gene trees. Overall, the accuracy is much poorer for increased ILS, as expected.

**Figure 5:**
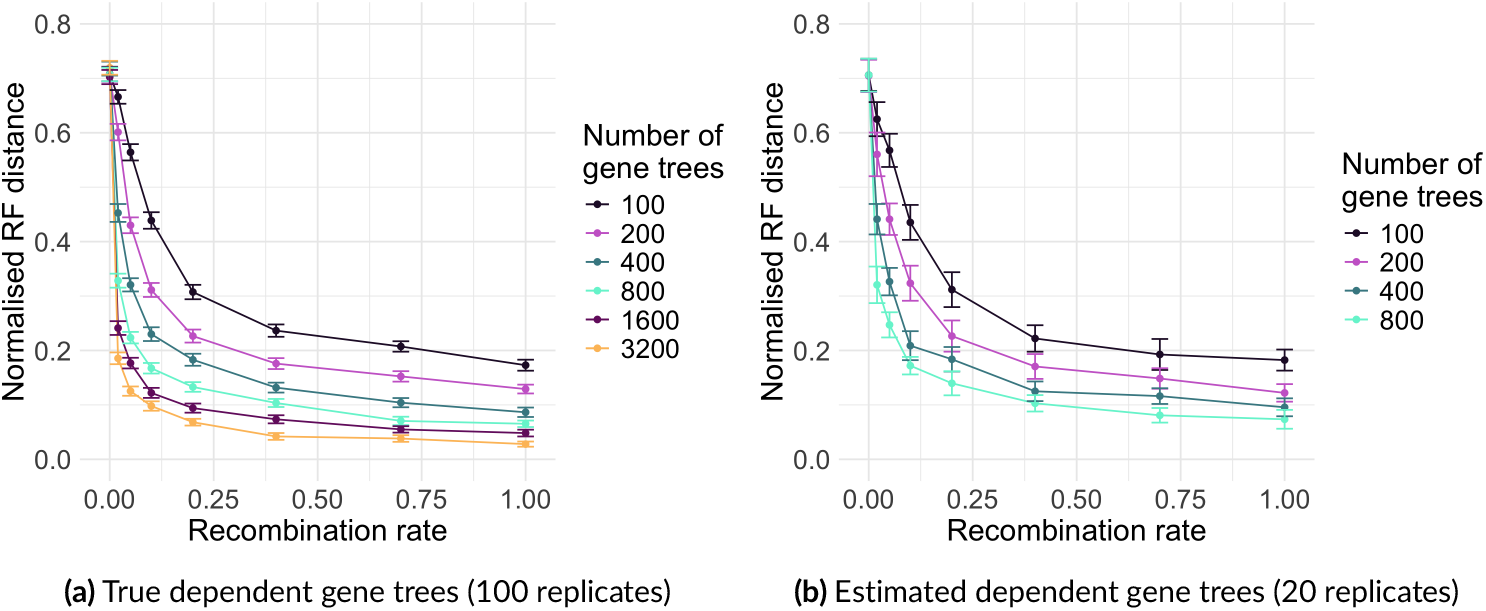
The accuracy of ASTRAL vs recombination rate with increased ILS (**0.2** ×). Error bars represent standard error.

### Different amounts of ILS

We also study the effect of different amounts of ILS on the accuracy of ASTRAL when gene trees are dependent. In Figure 6, we fix the number of gene trees to 200 and vary the branch length multiplier. We observe similar patterns of normalised RF distance with respect to recombination rate as above. Overall, ASTRAL performs better with less ILS, as observed for independent gene trees in Mirarab et al. (2014b).

**Figure 6:**
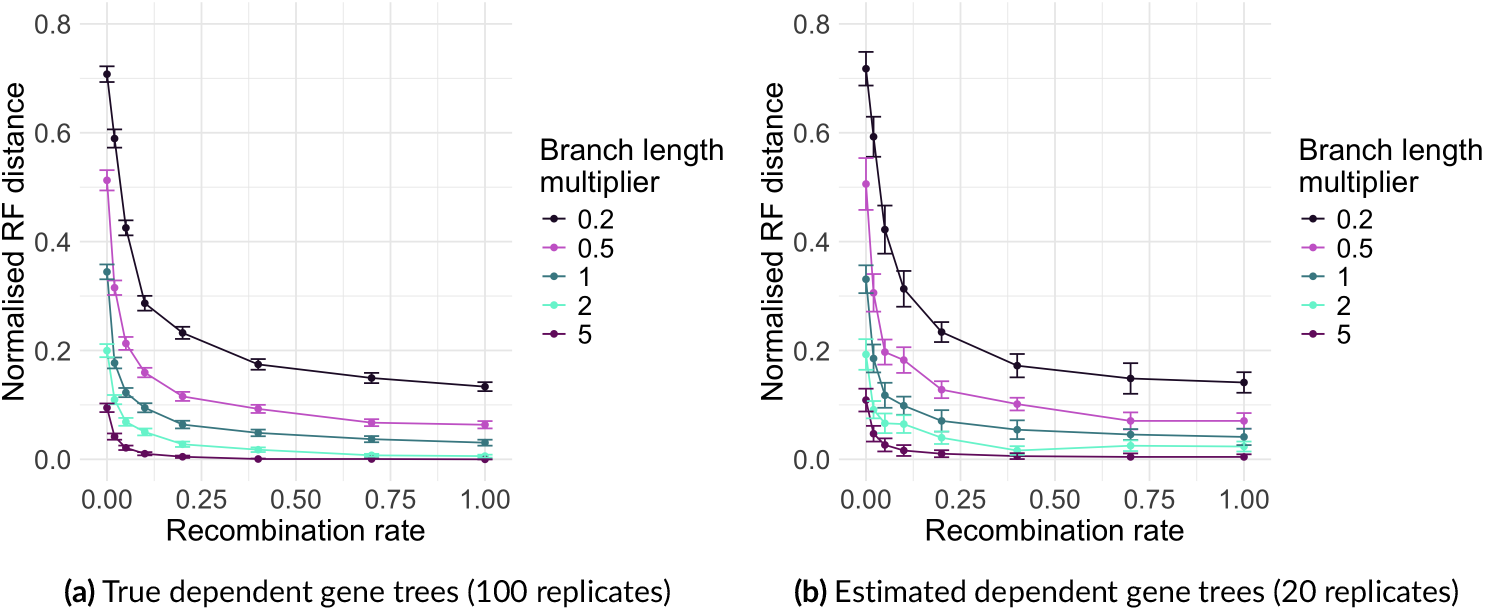
The accuracy of ASTRAL vs recombination rate with different amounts of ILS for **N = 200**. Error bars represent standard error.

### Varying recombination along the genome

We observe the effect of varying recombination along the genome by generating independent ‘blocks’ as described at the end of the Generating dependent gene trees section. We quantify the variability of the recombination rate by the proportion of independent gene trees (i.e., the proportion of gene trees that start a new block, which is the number of blocks divided by *N*). The results for fixed *R* (and therefore varying overall recombination rate) are shown in Figure 22 in Supporting information. As expected, we observe that ASTRAL becomes more accurate as the proportion of independent trees (and thus the overall recombination rate) increases.

In Figure 7 (see also Figure 23 in Supporting information), we fix the overall recombination rate to be roughly equivalent to the uniform dependence case with *R* = 0.1. We observe that as the number of blocks increases, the accuracy of ASTRAL degrades slightly; this effect is more noticeable for smaller numbers of gene trees. Thus more dependence within smaller blocks (i.e., more variation in recombination rate along the genome) is slightly worse than less dependence within larger blocks. At some point, the recombination rate within blocks becomes 0, and the trees within each block are forced to be identical; this is equivalent to having a smaller number of completely independent trees, and results in the lowest accuracy.

**Figure 7:**
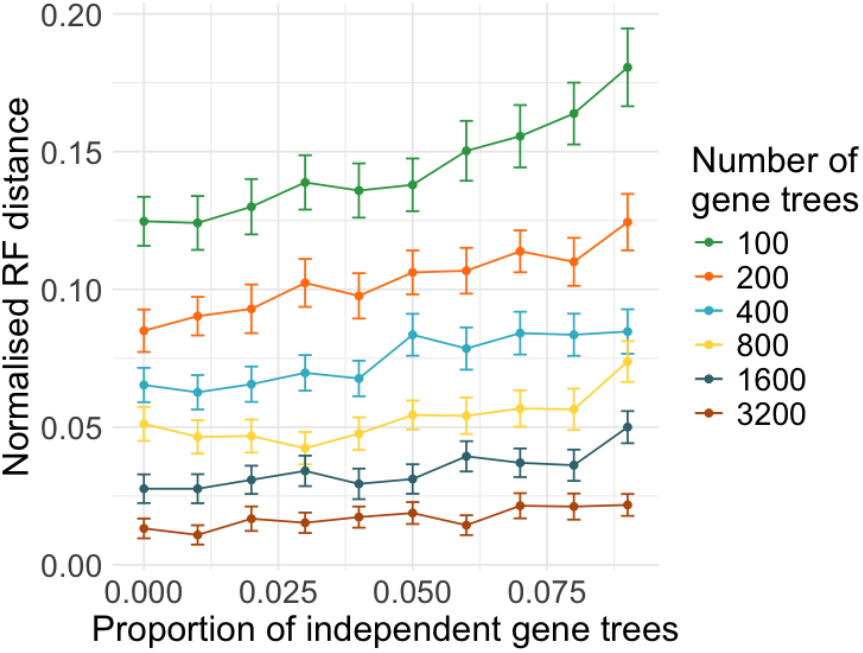
The accuracy of ASTRAL vs the proportion of independent gene trees. The accuracy is averaged over 100 replicates, with fixed overall recombination rate. Error bars represent standard error.

### Decreasing dependence by thinning trees

ASTRAL is constructed under the assumption that the gene trees are independent, and our results above show that it is more accurate in this case. It is common practice to ensure that this is the case by ‘thinning’ the gene trees, i.e., only considering trees that are a certain distance apart in the genome (Mongiardino Koch, 2021; Patané et al., 2024; Simmons et al., 2016). If this distance is chosen to be sufficiently large, the resulting trees can be considered to be approximately independent. On the other hand, thinning the trees in this way will reduce their number, which will also degrade the performance of ASTRAL.

To study this effect, we simulate 1000 dependent gene trees from the 37-taxon mammalian species tree for several recombination rates (0.02, 0.1, and 1). We then thin our dataset by subsampling every *n*th tree, for *n* ranging from 1 to 50, and use the resulting trees as input into ASTRAL. This is replicated 1000 times, with the results shown in Figure 8.

**Figure 8:**
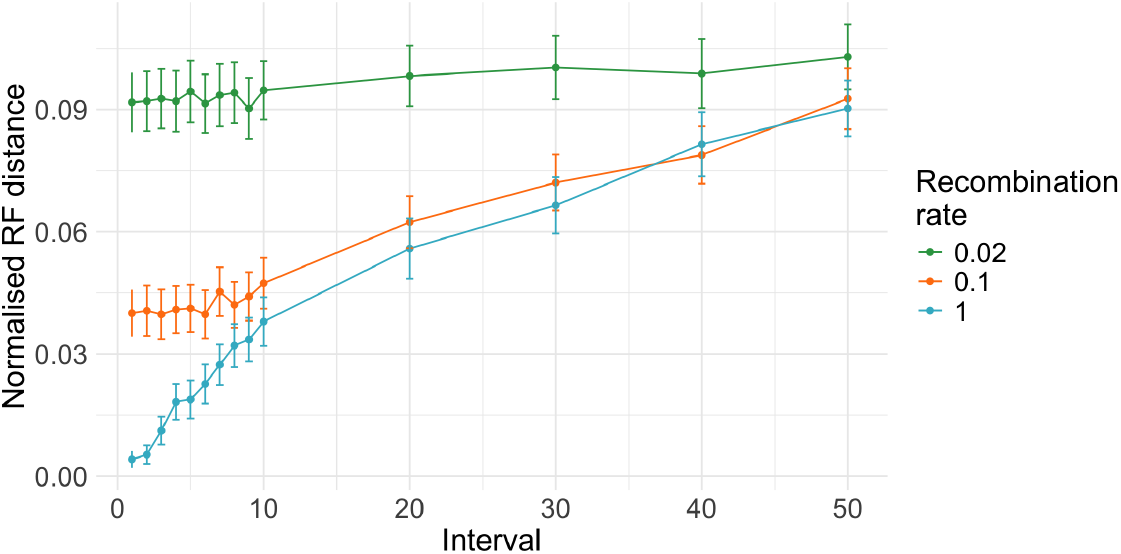
Accuracy of ASTRAL when thinning dependent gene trees for the 37-taxon mammalian species tree over 1000 replicates. Error bars represent standard error.

From this Figure, it is clear that using the full gene tree set (no thinning) is always the most accurate; accuracy decreases monotonically as the subsampling interval increases. Thus, it is always preferable to use a larger number of gene trees, even if they are dependent, over a smaller number of independent gene trees.

### Effective sample size

The multiplicative decrease in normalised RF distance due to gene tree dependence varies with the number of gene trees for fixed recombination rate, so it does not provide an overall measure of the effect of gene tree dependence. In order to obtain such a measure, we wish to quantify the amount of information or statistical power available in a sample with gene tree dependence; in other words, the effective sample size (*ESS*). To determine the *ESS*, we first calculate the average normalised RF distance for ASTRAL for a recombination rate *R*. We then use a binary search to determine the number of independent gene trees required to produce the same normalised RF distance.

We can repeat this procedure for any sample size to fit a relationship between actual and effective sample size for any recombination rate. As an illustrative example, we consider *R* = 0.1. The results are shown in Figure 9, where the relationship between *ESS* and *N* appears approximately linear. Fitting a regression line through the origin yields

**Figure 9:**
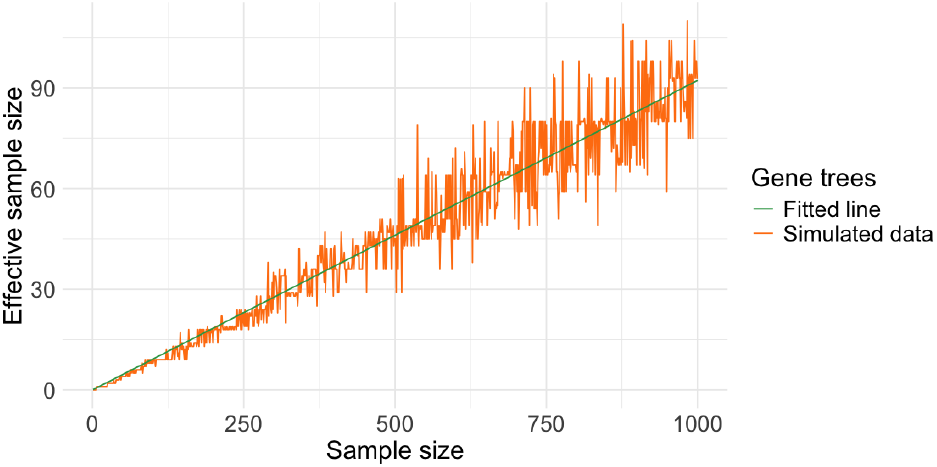
**The effective sample size of ASTRAL under dependence for** *R* = 0.1.

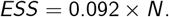

This quantifies the effect of gene tree dependence for this recombination rate, for any sample size. We further examine how the effective sample size varies with recombination rate. Figure 10 shows that *ESS* approaches the actual sample size *N* as the recombination rate increases, as expected, reflecting a reduction in gene tree dependence.

**Figure 10:**
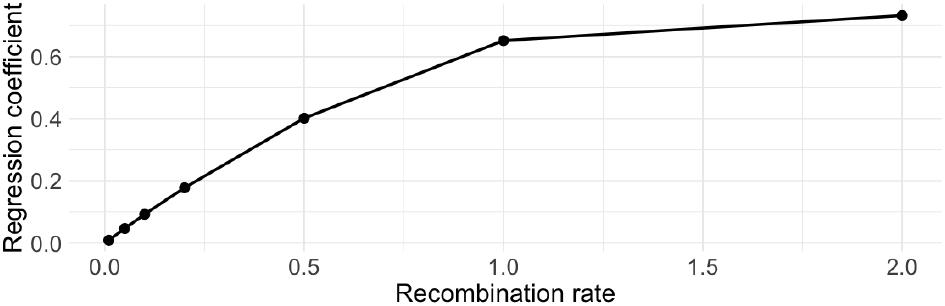
Regression coe”cient *α* as a function of recombination rate *R*. The coefficient *α* is obtained from the linear relationship *ESS* = *αN*.

### Robustness of our results

*16-taxon fungal species tree*. In Figure 24 in Supporting information, we observe similar patterns to those observed for the 37-taxon mammalian species tree, but with overall higher accuracy. This indicates that our results are not specific to the mammalian species tree. However, the numerical results could differ depending on the specific target phylogeny.

*Comparison with msprime*. Figures 24 and 25 in Supporting information show the accuracy of ASTRAL with dependent gene trees generated by our model and `msprime` for the 16-taxon fungal tree and for the 37-taxon mammalian tree respectively. It can be seen that although `msprime` results in a slightly lower accuracy, the results for the two models are in general very similar (the average absolute difference between the mean normalised RF distance of the two models is 0.0083 and 0.0098 for the mammlian and fungal trees respectively).

### Adjacent genes

To compare our numerical results with Wang and Liu (2016), we use our model on their 9-taxon mouse species tree, reproduced in Figure 26 in Supporting information. Because they divide the simulated genome into genes by recognising breakpoints, their genes are separated by a single recombination. To calibrate the corresponding recombination rate *R*, we calculate the average number of recombinations between consecutive gene trees in our model (from 10000 replicates), and use a binary search so that this average is 1. By repeating this process 20 times, we estimate an equivalent recombination rate of *R* = 0.09 (standard error 0.0001) for this tree.

We then fix the number of gene trees to the average produced by `ms` for the same parameters as Wang and Liu (2016); for the population recombination rates *ε* = 100, 200, 1000 that they used, this results in 867, 1752, and 8739 gene trees respectively. We then perform simulations as above for gene trees that are dependent and independent. To estimate the effect of gene tree estimation error, we also evolve sequences from true gene trees under the HKY85+Г (Hasegawa et al., 1985) model with *α* = 0.8, as in Wang and Liu (2016), and then estimate gene trees with FastTree (Price et al., 2009, 2010). Due to computational constraints, we run 100 replicates for inferred gene trees and 1000 replicates for true gene trees.

In Figure 11, we observe that under these parameters, the effect of gene tree dependence on ASTRAL is large for the smaller recombination rates, and substantially greater than that of gene tree estimation error. In fact, when the gene trees are independent, the species tree is inferred without error. This suggests that gene tree dependence can have a pronounced effect when genes are positioned directly adjacent to each other on the chromosome.

**Figure 11:**
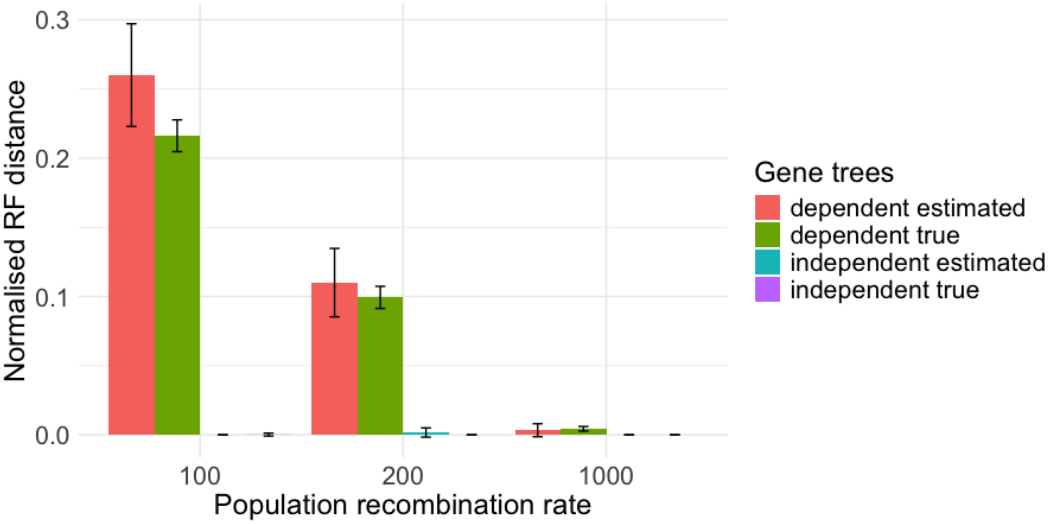
The accuracy of ASTRAL for the empirical tree of Wang and Liu (2016) for recombination rate R = 0.09. Error bars represent standard error.

Our results agree with Wang and Liu (2016) in showing a large effect of gene tree dependence. We do note that the numerical accuracies we obtain are slightly different from the equivalent cases in Wang and Liu (2016); apart from the differences between the models themselves, we also fix the recombination rate, rather than the number of recombinations between gene trees, and fix the number of gene trees. However, because the genes are taken to be directly adjacent to each other, this scenario overestimates the amount of dependence in a practical genomic dataset.

## Real data experiments

### Estimating minimum genomic distances for a SNP dataset

The accuracy of ASTRAL changes little when the recombination rate *R* between genes is larger than 1. With typical human-like parameters of *N*_*e*_ = 10^4^ and *r* = 10^−8^, this corresponds to a genomic distance of approximately 5Kb between loci. This suggests that when genes are separated by at least this distance, the effect of gene tree dependence on ASTRAL will be minimal. However, studies of the human genome indicate that LD can extend much further, sometimes even beyond 100 Kb (Abecasis et al., 2001; Collins et al., 1999; Taillon-Miller et al., 2000). It is common to use an LD-based threshold to extract putatively independent genes; however, as illustrated in Section Decreasing dependence by thinning trees, removing dependent genes can potentially lower inference accuracy. It would be preferable to establish a criterion whereby slightly dependent genes can still be used, as long as inference accuracy is minimally affected.

In order to do this, we re-analysed the biological data from Wang and Liu (2016). Here, SNP data from house mice (Liu et al., 2015) were analyzed, covering 19 chromosomes with an average of 21,085 SNPs per chromosome. For each chromosome, LD decay plots were used to determine a safe distance between concatenated SNPs, with distances determined to be betwen 1–3Mb.

In order to determine a safe distance using our methodology, we used their inferred species tree and simulated gene trees under our model, varying both the number of genes (30, 50, 70) and the recombination rate. We then observed the normalised RF distance between the inferred and original species tree, comparing it to the distance produced when using independent gene trees as a baseline.

The recombination rate *R* between genes is related linearly to genomic distance via population size and per-site recombination rate. Because the effective population size *N*_*e*_ varies across studies, we adopt a conservative lower-bound value of 10^4^ (Deinum et al., 2015; Geraldes et al., 2008; Phifer-Rixey et al., 2020). With a recombination rate per site per generation of *r* = 10^−8^, this gives a relation of *d* = 5000*R*.

Figure 12 shows the increase in normalised RF distance from the independent to the dependent case, with the difference approaching zero as *d* increases. When the difference becomes negligible, it is safe to use genes that are separated by this distance, since longer distance yields little gain in accuracy. To account for stochastic variation, we define the threshold as approximately twice the standard error of the accuracy from independent gene trees (0.05). This results in different cut-off values across various number of genes, as shown in Table 2. However, all the genomic distances we identified are orders of magnitude smaller (in kilobases rather than megabases) than those required under LD-based methods (Wang and Liu, 2016). This suggests that we can use relatively close loci to achieve a better accuracy for species tree inference.

**Table 2:** Cut-off values (Kb) of genomic distance between genes.

| Number of genes | Minimum distance |
| --- | --- |
| 30 | 2.5 |
| 50 | 3.5 |
| 70 | 1 |

**Figure 12:**
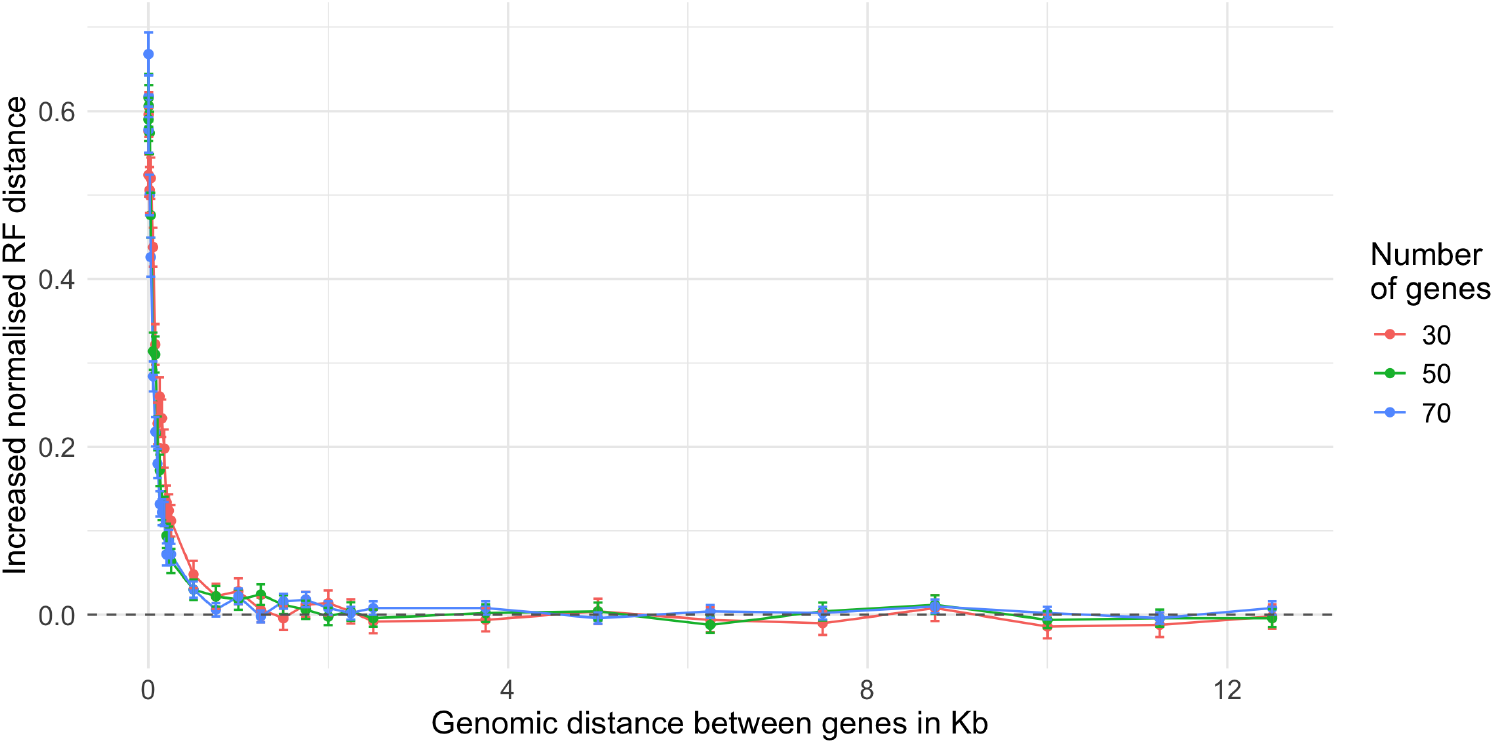
Decrease of accuracy of ASTRAL from using independent to dependent gene trees vs genomic distance between loci, averaged over 100 replicates. Error bars represent standard error.

This also shows a procedure by which we can estimate these ‘safe’ distances; although it is more complicated than an LD-based calculation and requires a model tree and some estimated parameters, this allows us to justifiably select loci that are close together without too much concern about whether they are truly independent.

### Estimating minimum genomic distance in a phylogenomic dataset

Our method can also be applied to datasets where orthologous genes have been identified and extracted from whole genome sequences. As an illustration, we re-analyse the biological gene trees from Song et al. (2012) on the 37-taxon mammalian species tree studied above. This dataset contains 424 effective genes (after 21 genes with mislabelled sequences and 2 outliers were removed from the original 447-gene dataset (Mirarab et al., 2014b)) with average length 3157bp. We re-estimated the gene trees with IQ-TREE (Minh et al., 2020; Nguyen et al., 2015).

Although the order of genes is not necessarily conserved between species, we located the position of each gene in the well-studied human genome. The average genomic distance between genes in this dataset is *d* = 5,817,377bp, with a corresponding recombination rate between gene trees of approximately 1163.48 (using *N*_*e*_ = 10^4^, *r* = 10^−8^). This is far above the range at which dependence affects the accuracy of ASTRAL, suggesting that dependence has little effect in this dataset. To check this, we compute the average pairwise normalised RF distance between gene trees separated by a fixed number of intervening genes. In Figure 13, we see that genes that are immediately consecutive in the dataset exhibit a slightly lower RF distance, indicating a small dependence, but this effect vanishes for genes that are even 2–3 genes apart.

**Figure 13:**
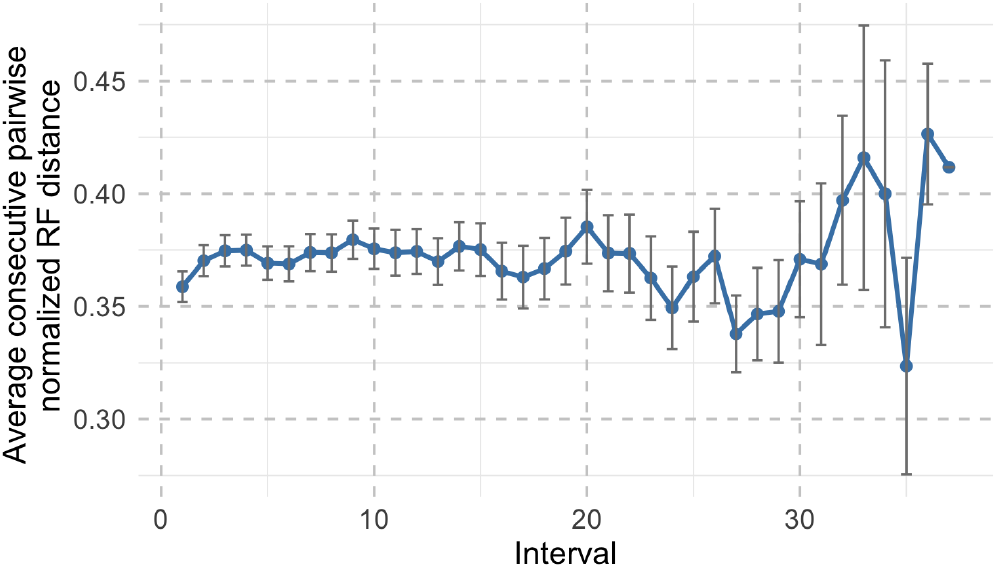
The average normalised RF distance between gene trees that are separated by a certain number of genes in the Song et al. (2012) dataset. Error bars represent standard error.

However, this dataset is an older dataset containing only a few hundred genes. More recent mammalian datasets (e.g., (Allio et al., 2024)) include thousands of genes, which greatly decreases the genomic distance between them. This decrease could enhance dependence between genes and potentially affect the accuracy of ASTRAL in species tree inference. By using the method proposed in Section Estimating minimum genomic distances for a SNP dataset, we can assess whether the genomic distances between genes are sufficiently large to ensure a minimal effect.

## Discussion

In this paper, we developed a model to simulate dependent gene trees within a species tree under a realistic process of incomplete lineage sorting and recombination. We then reassessed the accuracy of ASTRAL for when gene trees are dependent, including both true and estimated gene trees. Our results show that ASTRAL performs more poorly as the dependence between gene trees increases, and gene tree estimation error adversely affects the accuracy of ASTRAL under dependence. Our simulations can quantify this effect for any recombination distance between genes.

Our findings are particularly relevant as whole-genome sequencing produces more gene trees for each genome. It is tempting to think that increasing the number of gene trees will counter the inaccuracy resulting from gene tree dependence. However, the number of genes in a genome is not unlimited, and furthermore, including more gene trees will lower the genomic distance between the trees, and thus increase the dependence between them, making the effect of gene tree dependence more pronounced. Thus, as more and more data is collected, it becomes more important to consider the effect of gene tree dependence. For the relatively small species trees that we study here, we can achieve a satisfactory reconstruction accuracy with relatively few gene trees even when dependence is taken into account, but this will not necessarily always be the case. For larger trees that are more difficult to reconstruct exactly, the number of gene trees needed to achieve a satisfactory accuracy may well exceed the actual number of genes available.

In our re-analysis of real datasets, we find that extracting orthologous genes from wholegenome sequences tends to produce genes that are sufficiently far apart that dependence has minimal effect. Conversely, on a SNP dataset, dependence can be significant if loci are taken to be directly adjacent. While LD-based thresholds are used to ensure independent loci, our findings indicate that they are too strict: achieving full independence between genes is unnecessary, and may even reduce accuracy by substantially limiting the number of informative loci available for analysis. Our results show a way by which we can estimate a safe distance between loci.

Although we have restricted our simulations to ASTRAL as it is currently the most popular summary method for species tree inference, it is likely that many other summary methods (Han and Molloy, 2023, 2025; Mahbub et al., 2021; Zhang and Mirarab, 2022a,b; Zhang et al., 2020) (in particular, methods that attempt to solve the maximum quartet support problem as ASTRAL does) will be similarly affected by gene tree dependence. Virtually all simulations evaluating these methods are performed with independent gene trees, and therefore our results are applicable in these cases as well.

While our analysis of the SNPs data provides valuable insights into the impact of genomic distance between loci on the accuracy of ASTRAL, several limitations should be noted. We adopted a fixed effective population size *N*_*e*_ to convert recombination rates to genomic distances, but variation in *N*_*e*_ among species and through time could shift the inferred thresholds. We have used a conservative value that maximises the inferred ‘safe’ distance. Moreover, our empirical study focuses on the house mouse, and extrapolation to other species should be made cautiously.

When estimating the the effective sample size for ASTRAL, there is an accumulated error including the accuracy of different sample sizes, and the matching of sample sizes to a given accuracy. Future work includes proposing a simpler and more reliable method to estimate the effective sample size. In addition, estimating the effective sample size for a dataset requires first estimating the recombination rate between gene trees, which introduces further uncertainty.

Our results suggest that a new species tree inference method to take gene tree dependence into account should be designed. One possible way is to assign different weights to quartets according to their patterns of occurrence in the genome. By incorporating the information of genomic location, this new method could potentially be more accurate in the case of gene tree dependence. This is the subject of future work.

## Data and code availability

Data and code are available on https://github.com/WAHE1/ASTI-GTD/tree/main.

## Supporting information

### Re-estimating branch lengths for the 37-taxon mammalian tree

The terminal branch lengths of the species tree given in Mirarab et al. (2014b) are all 9.0 coalescent units long; this is an artifact of the MP-EST summary method used to estimate it. This is immaterial for the topologies of the gene trees simulated under the MSC from this species tree; however, it has an effect on the branch lengths of these trees, and therefore on sequences generated from these trees.

To amend this, we re-estimate the branch lengths for the 37-taxon mammalian tree from the original data of Song et al. (2012), keeping the topology of the tree used in Mirarab et al. (2014b). We compute a distance matrix of the 447 genes from the gene trees, and use them as input to ERaBLE (Binet et al., 2016), which also estimates the relative evolutionary rates of the genes. The resulting tree is then re-rooted on the leaf branch ‘Gal’ by splitting the branch into two halves so that the length of the branch to ‘Gal’ is the average of the lengths of the paths from the root to the other leaf nodes.

The branch lengths of the resulting tree are expressed in units of (average number of) substitutions per site. To translate this into coalescent units (in the absence of estimated mutation rates and population sizes), we scale the tree by a constant factor so that the average (over all branch lengths *l*) of 1 – *e*^*-l*^ is equal to that of the tree in Mirarab et al. (2014b). This can be thought of as an approximate ‘amount of coalescence’ in the tree. We also need to convert simulated gene trees back into substitutions per site to evolve sequences along them. This is achieved by rescaling the branch lengths using the inverse of the same scaling factor. In addition, we sample a rate inferred by ERaBLE from a randomly selected gene tree and use this rate to further rescale the branch lengths.

To verify that our gene trees have more realistic topology and branch lengths than the trees from Mirarab et al. (2014b), we simulated 100 gene trees from the MSC model on our species tree. We then calculated the pairwise normalised RF distance (Robinson and Foulds, 1981) and the Kuhner–Felsenstein (KF, aka branch score) distance (Kuhner and Felsenstein, 1994) between all simulated gene trees, biological gene trees from Song et al. (2012), and simulated gene trees from Mirarab et al. (2014b). These distances were used to produce a Multidimensional Scaling (MDS) plot (Figure 14), which shows that our simulated gene trees more closely resemble the biological trees.

### Parameters for gene evolution

According to Mirarab et al. (2014a), the parameters used for evolving sequences in Section Simulations, listed below, were estimated from avian data (Jarvis et al., 2014) by the program bppml, also from bppsuite (Guéguen et al., 2013). Mirarab et al. (2014b) ran bppml on the subset of 1185 avian genes that included all the taxa.

- Substitution model parameters:
  – a=1.062409952497,
  – b=0.133307705766,
  – c=0.195517800882,
  – d=0.223514845018,
  – e=0.294405416545,
  – theta=0.469075709819,
  – theta1=0.558949940165,
  – theta2=0.488093447144

- Rate distribution parameters for Gamma:
  – n=4,
  – alpha=0.370209777709

## Supplementary figures

**Figure 14:**
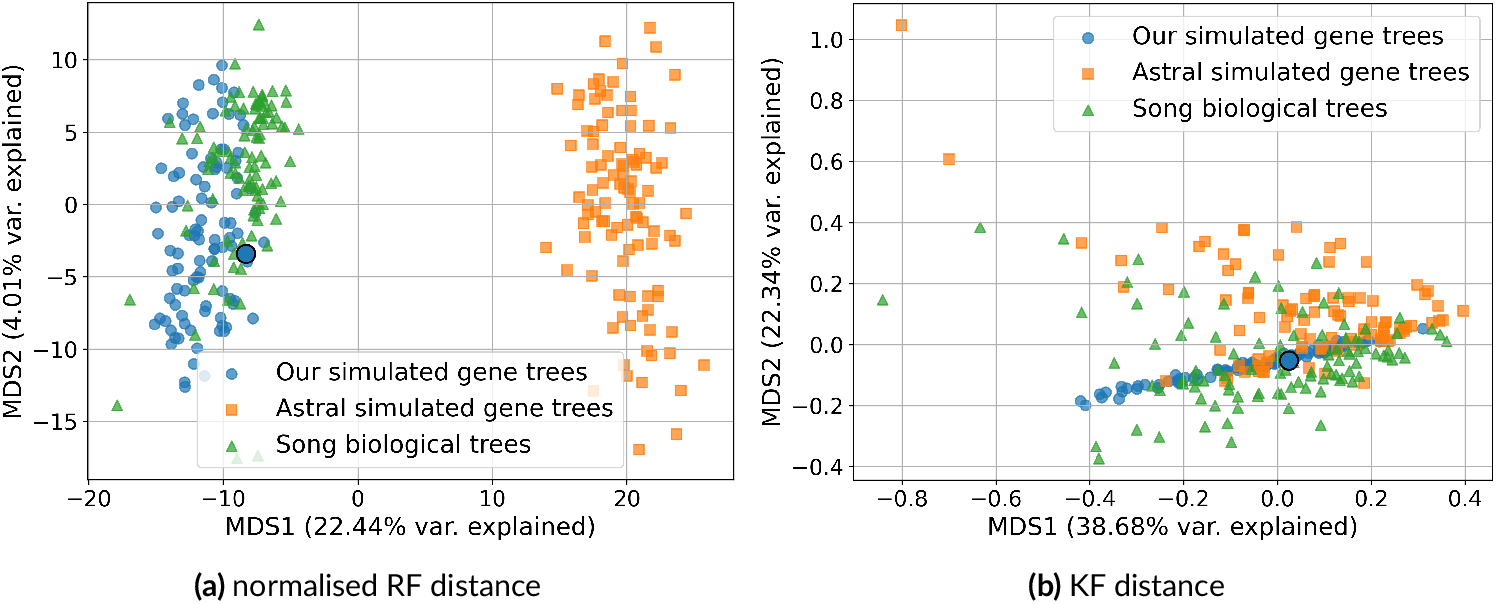
MDS analysis of simulated gene trees and biological trees. The blue circles represent our species tree.

**Figure 15:**
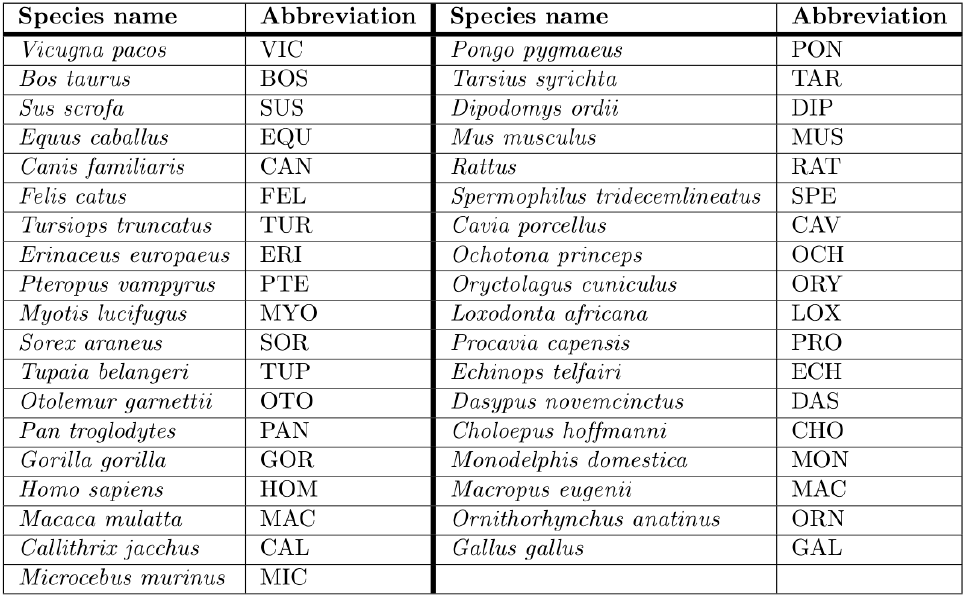
Abbreviations of species names in the 37-taxon mammalian species tree in Fig 3.

**Figure 16:**
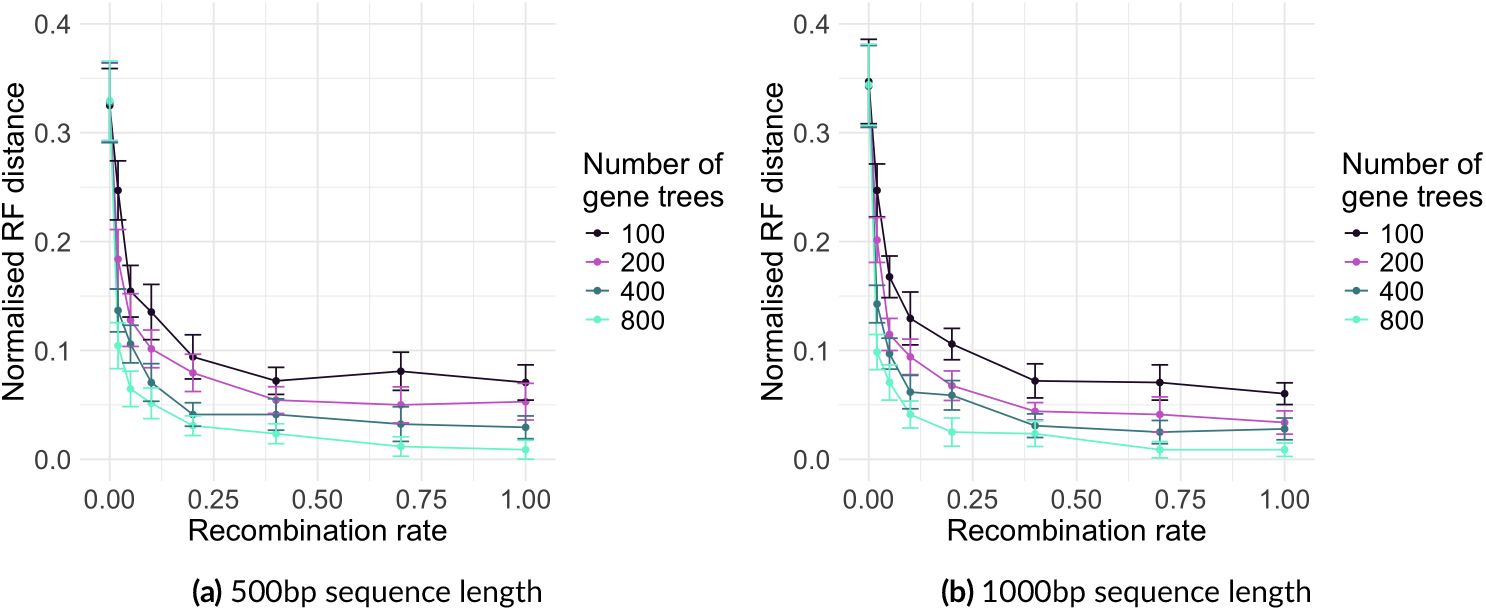
The accuracy of ASTRAL vs recombination rate for estimated gene trees, averaged over 20 replicates. Error bars represent standard error.

**Figure 17:**
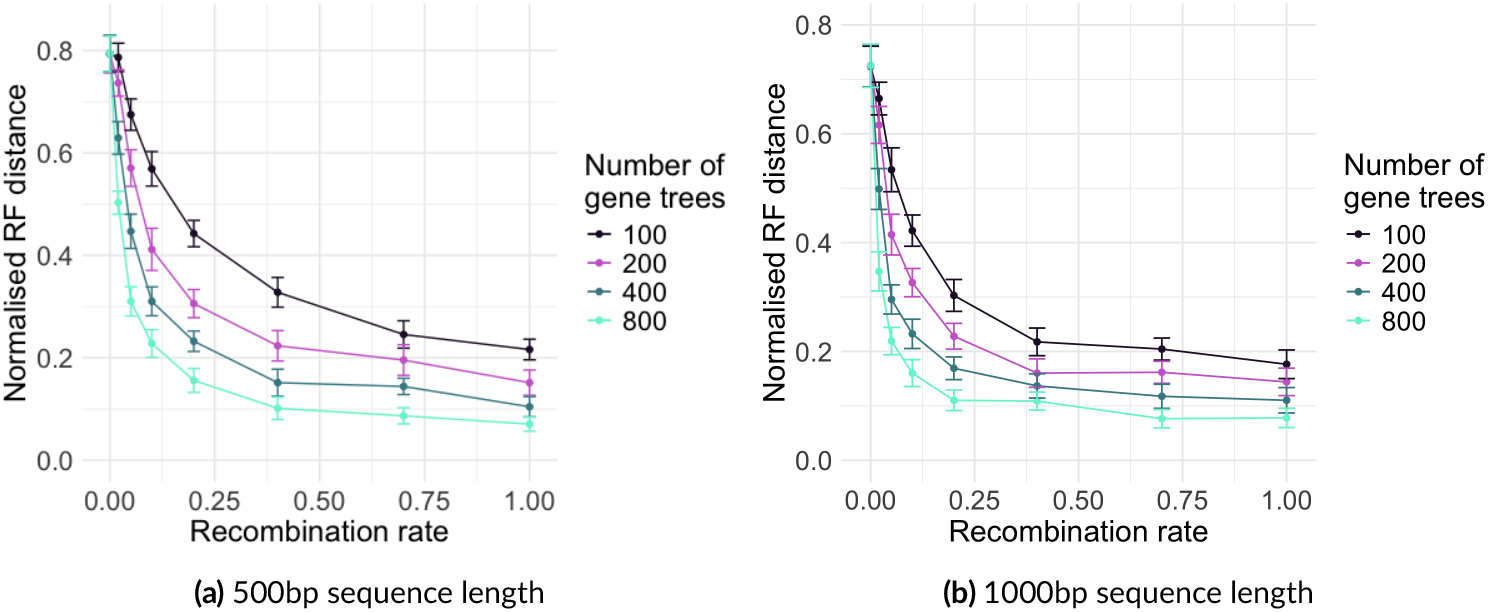
The accuracy of ASTRAL vs recombination rate with increased ILS (0.2×) for estimated gene trees, averaged over 20 replicates. Error bars represent standard error.

**Figure 18:**
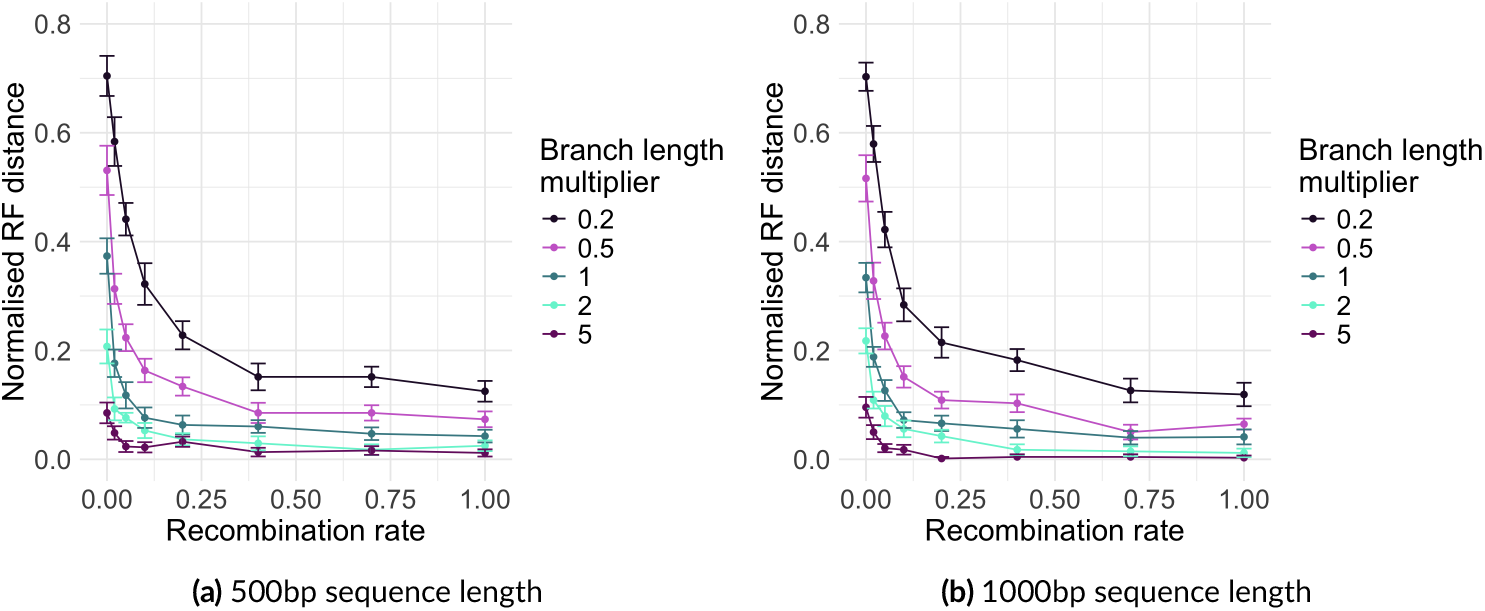
The accuracy of ASTRAL vs recombination rate with different amount of ILS for estimated gene trees, averaged over 20 replicates. Error bars represent standard error.

**Figure 19:**
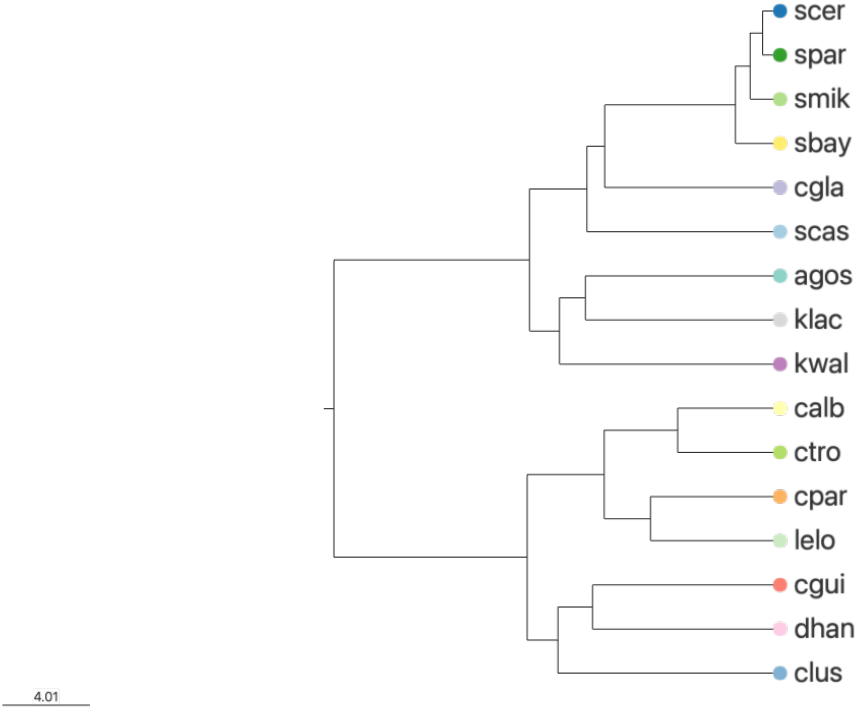
The 16-taxon fungi species tree with branch lengths in coalescent units. Species names are given in Table 20 in Supporting information.

**Figure 20:**
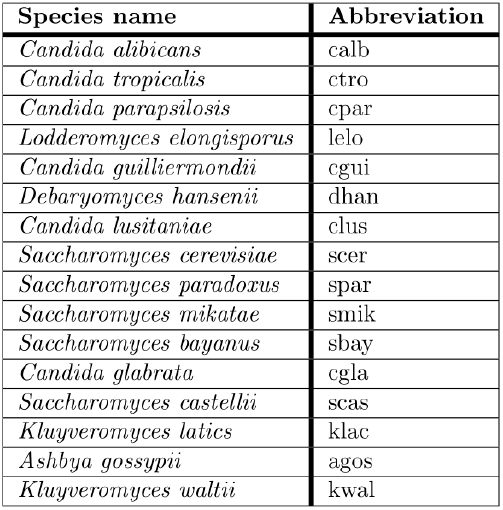
Abbreviations of species names in the 16-taxon fungi species tree in Figure 19 in Supporting information.

**Figure 21:**
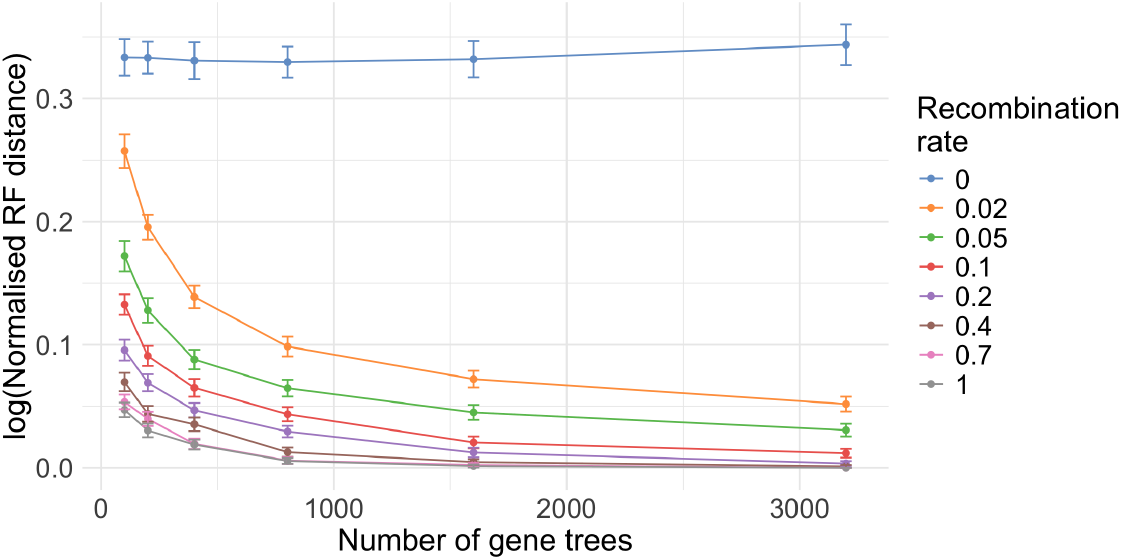
The accuracy of ASTRAL (log normalised RF distance) vs the number of gene trees, averaged over 1000 replicates.

**Figure 22:**
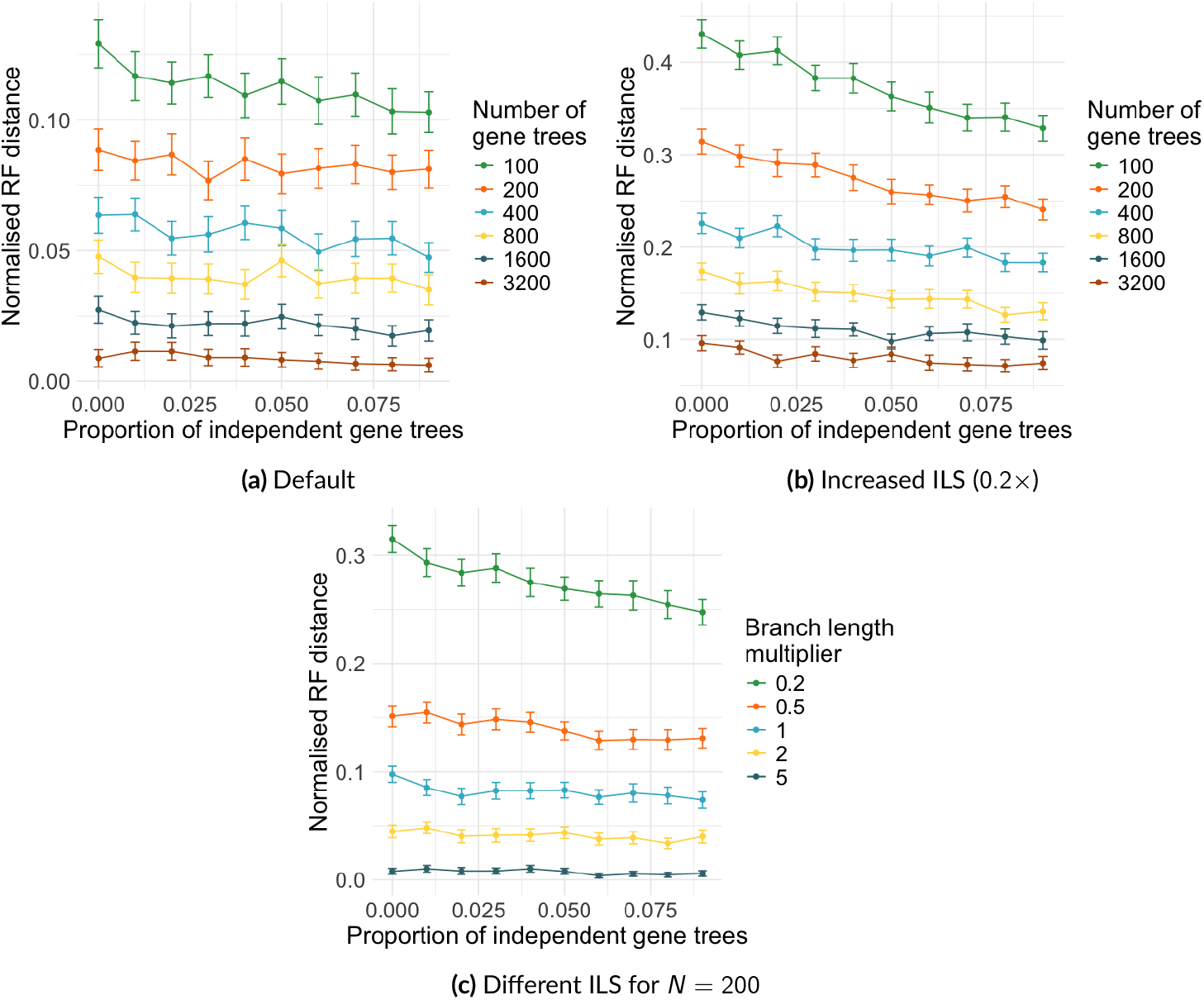
The accuracy of ASTRAL vs the proportion of independent gene trees, averaged over 100 replicates, with block-fixed recombination rate of 0.1. Error bars represent standard error.

**Figure 23:**
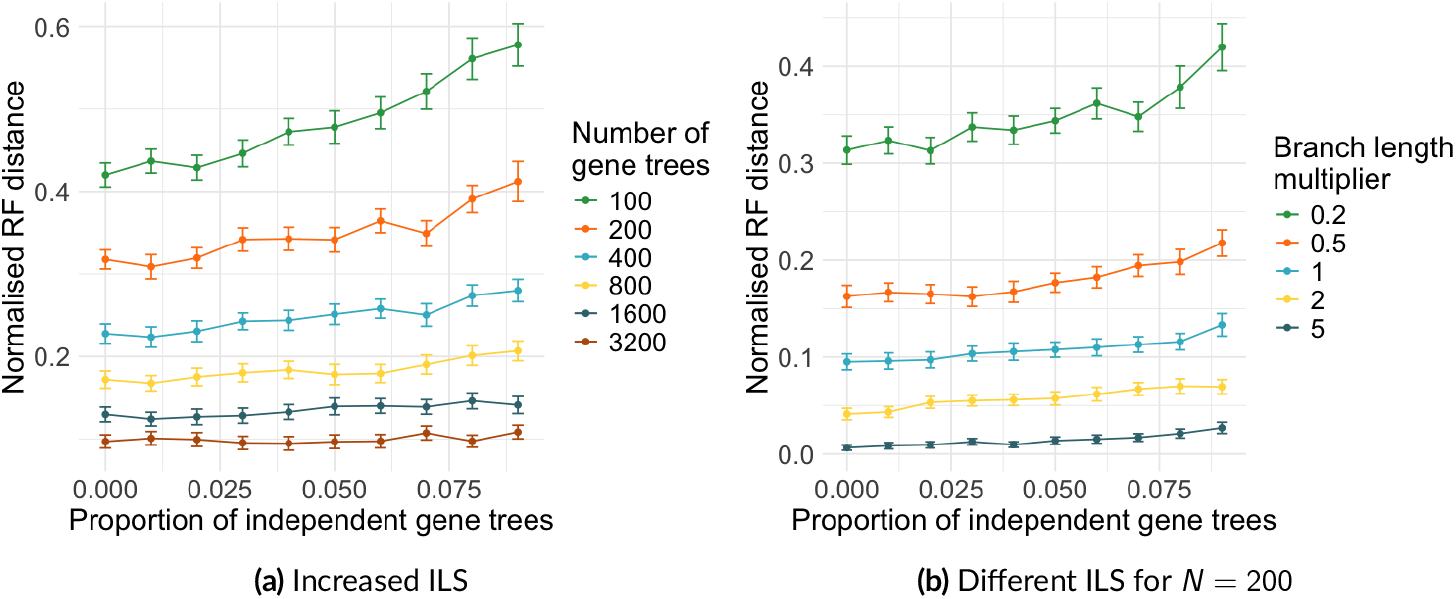
The accuracy of ASTRAL vs the proportion of independent gene trees, averaged over 100 replicates, with fixed overall recombination rate of 0.1. Error bars represent standard error. The default settings are shown in Figure 7.

**Figure 24:**
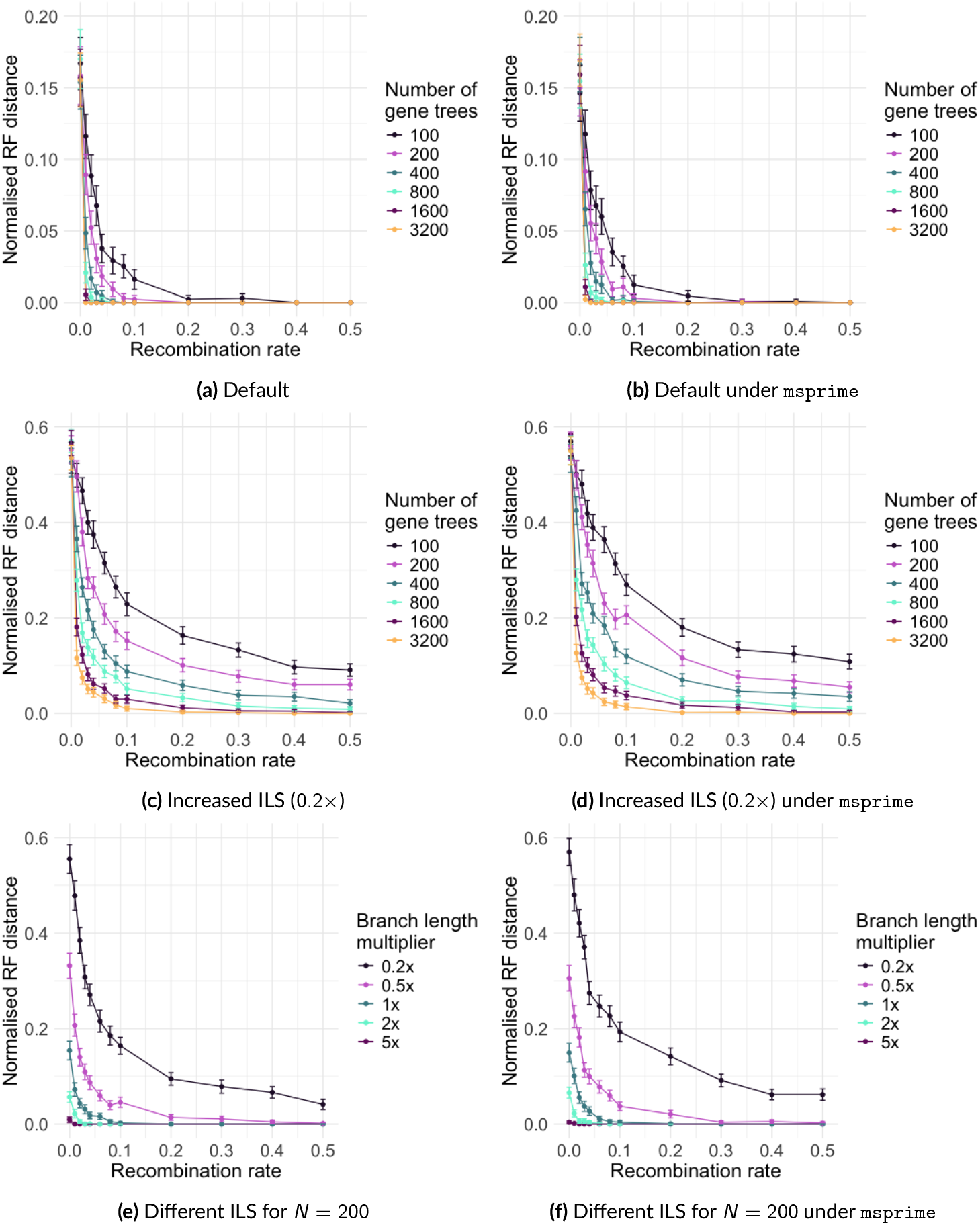
The accuracy of ASTRAL vs recombination rate for the 16-taxon fungi species tree, averaged over 100 replicates. Error bars represent standard error.

**Figure 25:**
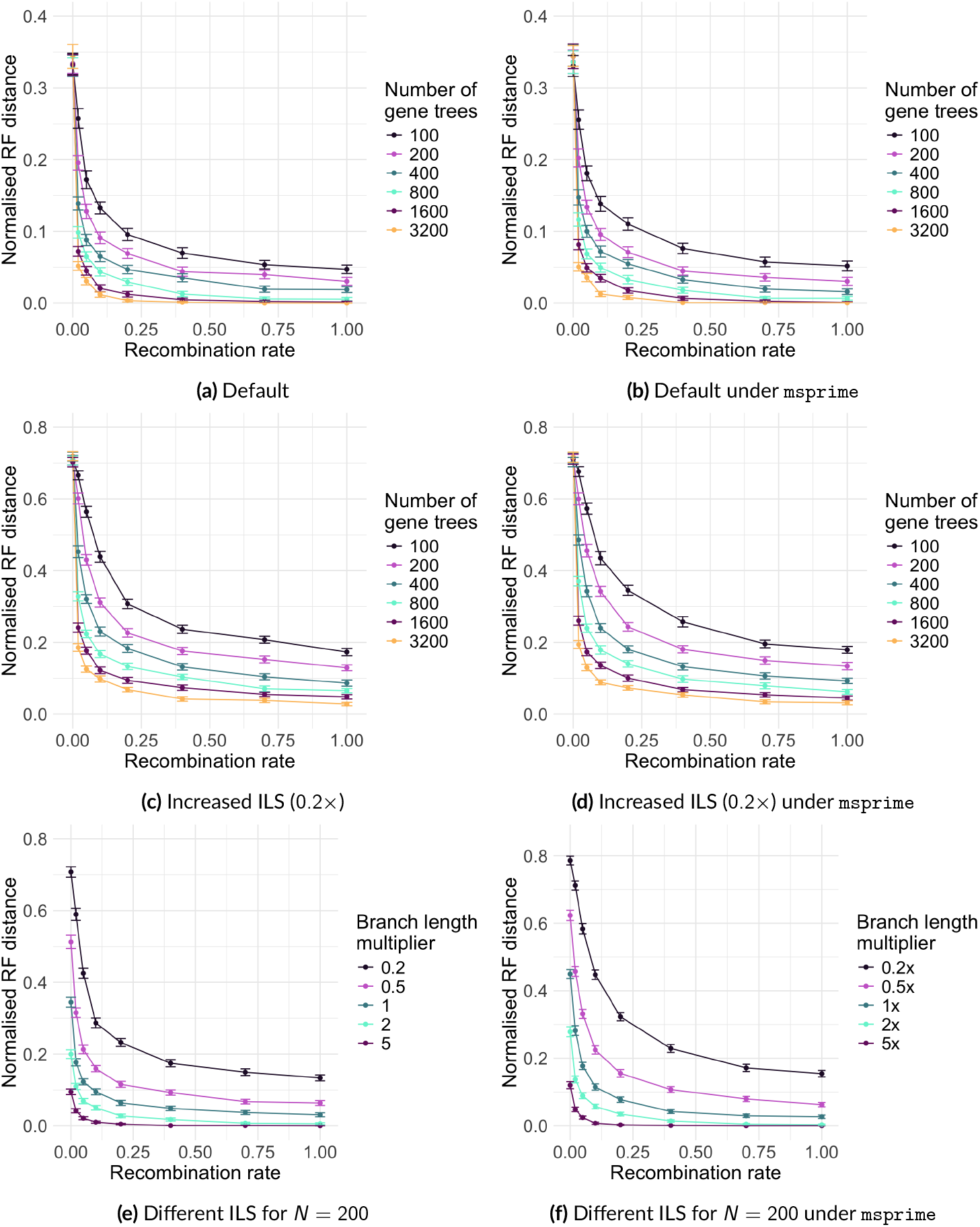
The accuracy of ASTRAL vs recombination rate for the 37-taxon mammalian species tree, averaged over 100 replicates. Error bars represent standard error.

**Figure 26:**
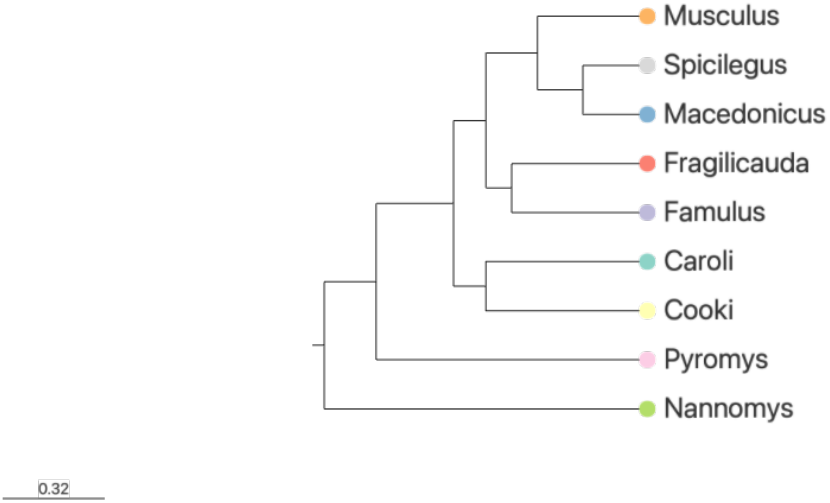
The 9-taxon mouse species tree from Wang and Liu (2016).

## References

Abecasis GR, Noguchi E, Heinzmann A, Traherne JA, Bhattacharyya S, Leaves NI, Anderson GG, Zhang Y, Lench NJ, Carey A, et al. (2001). Extent and distribution of linkage disequilibrium in three genomic regions. The American Journal of Human Genetics 68, 191–197.

Allio R, Delsuc F, Belkhir K, Douzery EJ, Ranwez V, Scornavacca C (2024). OrthoMaM v12: A database of curated single-copy ortholog alignments and trees to study mammalian evolutionary genomics. Nucleic Acids Research 52, D529–D535.

Ballesteros JA, Sharma PP (2019). A critical appraisal of the placement of Xiphosura (Chelicerata) with account of known sources of phylogenetic error. Systematic Biology 68, 896–917.

Barker D, Pagel M (2005). Predicting Functional Gene Links from Phylogenetic-Statistical Analyses of Whole Genomes. PLoS Computational Biology 1, 1–8.

Baumdicker F, Bisschop G, Goldstein D, Gower G, Ragsdale AP, Tsambos G, Zhu S, Eldon B, Ellerman EC, Galloway JG, Gladstein AL, Gorjanc G, Guo B, Jeffery B, Kretzschumar WW, Lohse K, Matschiner M, Nelson D, Pope NS, Quinto-Cortés CD, et al. (2022). EPcient ancestry and mutation simulation with msprime 1.0. Genetics 220, iyab229.

Binet M, Gascuel O, Scornavacca C, P. Douzery EJ, Pardi F (2016). Fast and accurate branch lengths estimation for phylogenomic trees. BMC bioinformatics 17, 23.

Butler G, Rasmussen MD, Lin MF, Santos MA, Sakthikumar S, Munro CA, Rheinbay E, Grabherr M, Forche A, Reedy JL, et al. (2009). Evolution of pathogenicity and sexual reproduction in eight Candida genomes. Nature 459, 657–662.

Chan Y, Li Q, Scornavacca C (2022). The large-sample asymptotic behaviour of quartet-based summary methods for species tree inference. Journal of Mathematical Biology 85, 1–22.

Chen GK, Marjoram P, Wall JD (2009). Fast and fexible simulation of DNA sequence data. Genome research 19, 136–142.

Collins A, Lonjou C, Morton N (1999). Genetic epidemiology of single-nucleotide polymorphisms. Proceedings of the National Academy of Sciences 96, 15173–15177.

Conry M (2020). Determining the impact of recombination on phylogenetic inference. The Florida State University.

Coyne J, Orr H (2004). Speciation. Sunderland, MA 276, 854.

Darwin C (1859). On the Origin of Species by Means of Natural Selection, or the Preservation of Favoured Races in the Struggle for Life. London: John Murray.

Davidson R, Vachaspati P, Mirarab S, Warnow T (2015). Phylogenomic species tree estimation in the presence of incomplete lineage sorting and horizontal gene transfer. BMC Genomics 16, 1–12.

DeGiorgio M, Degnan JH (2014). Robustness to divergence time underestimation when inferring species trees from estimated gene trees. Systematic Biology 63, 66–82.

Deinum EE, Halligan DL, Ness RW, Zhang YH, Cong L, Zhang JX, Keightley PD (2015). Recent evolution in Rattus norvegicus is shaped by declining eFective population size. Molecular biology and evolution 32, 2547–2558.

Flouri T, Jiao X, Rannala B, Yang Z (2018). Species tree inference with BPP using genomic sequences and the multispecies coalescent. Molecular biology and evolution 35, 2585–2593.

Geraldes A, Basset P, Gibson B, Smith KL, Harr B, Yu HT, Bulatova N, Ziv Y, Nachman MW (2008). Inferring the history of speciation in house mice from autosomal, X-linked, Y-linked and mitochondrial genes. Molecular ecology 17, 5349–5363.

Giarla TC, Esselstyn JA (2015). The challenges of resolving a rapid, recent radiation: empirical and simulated phylogenomics of Philippine shrews. Systematic Biology 64, 727–740.

Griffiths RC, Marjoram P (1997). An ancestral recombination graph. Institute for Mathematics and its Applications 87, 257.

Guéguen L, Gaillard S, Boussau B, Gouy M, Groussin M, Rochette NC, Bigot T, Fournier D, Pouyet F, Cahais V, et al. (2013). Bio++: ePcient extensible libraries and tools for computational molecular evolution. Molecular biology and evolution 30, 1745–1750.

Han Y, Molloy EK (2023). Improving quartet graph construction for scalable and accurate species tree estimation from gene trees. Genome Research 33, 1042–1052.

Han Y, Molloy EK (2025). Improved robustness to gene tree incompleteness, estimation errors, and systematic homology errors with weighted TREE-QMC. Systematic Biology, syaf009.

Hasegawa M, Kishino H, Yano Ta (1985). Dating of the human-ape splitting by a molecular clock of mitochondrial DNA. Journal of Molecular Evolution 22, 160–174.

Heled J, Drummond AJ (2009). Bayesian inference of species trees from multilocus data. Molecular biology and evolution 27, 570–580.

Hillis DM, Moritz C, Mable BK (1996). Molecular systematics. Vol. 23. Sinauer.

Huang H, Knowles LL (2016). Unforeseen consequences of excluding missing data from next-generation sequences: simulation study of RAD sequences. Systematic Biology 65, 357–365.

Hudson RR (1983). Properties of a neutral allele model with intragenic recombination. Theoretical Population Biology 23, 183–201.

Hudson RR (2002). Generating samples under a Wright–Fisher neutral model of genetic variation. Bioinformatics 18, 337–338.

Jarvis E, Mirarab S, Aberer A, Houde P, Li C, Ho S, Faircloth B, Nabholz B, Howard J, Suh A, Weber C, Fonseca R, Alfaro-Nunez A, Narula N, Liu L, Burt D, Ellegren H, Edwards S, Stamatakis A, Mindell D, et al. (2014). Phylogenomic analyses data of the avian phylogenomics project. 10.5524/101041. URL: http://gigadb.org/dataset/101041.

Kelleher J, Etheridge AM, McVean G (2016). EPcient coalescent simulation and genealogical analysis for large sample sizes. PLoS Computational Biology 12, e1004842.

Kingman JF (1982). On the genealogy of large populations. Journal of Applied Probability 19, 27–43.

Kuhner MK, Felsenstein J (1994). A simulation comparison of phylogeny algorithms under equal and unequal evolutionary rates. Molecular biology and evolution 11, 459–468.

Lanier HC, Knowles LL (2012). Is recombination a problem for species-tree analyses? Systematic Biology 61, 691–701.

Lanier HC, Knowles LL (2015). Applying species-tree analyses to deep phylogenetic histories: challenges and potential suggested from a survey of empirical phylogenetic studies. Molecular Phylogenetics and Evolution 83, 191–199.

Legried B, Molloy EK, Warnow T, Roch S (2021). Polynomial-time statistical estimation of species trees under gene duplication and loss. Journal of Computational Biology 28, 452–468.

Li Q, Scornavacca C, Galtier N, Chan Y (2021). The multilocus multispecies coalescent: a fexible new model of gene family evolution. Systematic Biology 70, 822–837.

Liu KJ, Steinberg E, Yozzo A, Song Y, Kohn MH, Nakhleh L (2015). Interspecific introgressive origin of genomic diversity in the house mouse. Proceedings of the National Academy of Sciences 112, 196–201.

Liu L (2008). BEST: Bayesian estimation of species trees under the coalescent model. Bioinformatics 24, 2542–2543.

Liu L, Yu L (2011). Estimating species trees from unrooted gene trees. Systematic Biology 60, 661–667.

Liu L, Yu L, Edwards SV (2010). A maximum pseudo-likelihood approach for estimating species trees under the coalescent model. BMC Evolutionary Biology 10, 1–18.

Liu L, Yu L, Pearl DK, Edwards SV (2009). Estimating species phylogenies using coalescence times among sequences. Systematic Biology 58, 468–477.

Maddison WP (1997). Gene trees and species trees. Systematic Biology 46, 523–536.

Mahbub M, Wahab Z, Reaz R, Rahman MS, Bayzid MS (2021). wQFM: highly accurate genomescale species tree estimation from weighted quartets. Bioinformatics 37, 3734–3743.

Markin A, Eulenstein O (2021). Quartet-based inference is statistically consistent under the unified duplication-loss-coalescence model. Bioinformatics 37, 4064–4074.

McVean GA, Cardin NJ (2005). Approximating the coalescent with recombination. Philosophical Transactions of the Royal Society B: Biological Sciences 360, 1387–1393.

Minh BQ, Schmidt HA, Chernomor O, Schrempf D, Woodhams MD, Von Haeseler A, Lanfear R (2020). IQ-TREE 2: new models and ePcient methods for phylogenetic inference in the genomic era. Molecular Biology and Evolution 37, 1530–1534.

Mirarab S, Bayzid MS, Boussau B, Warnow T (2014a). Statistical binning enables an accurate coalescent-based estimation of the avian tree. Science 346, 1250463.

Mirarab S, Reaz R, Bayzid MS, Zimmermann T, Swenson MS, Warnow T (2014b). ASTRAL: genomescale coalescent-based species tree estimation. Bioinformatics 30, i541–i548.

Mirarab S, Warnow T (2015). ASTRAL-II: coalescent-based species tree estimation with many hundreds of taxa and thousands of genes. Bioinformatics 31, i44–i52.

Molloy EK, Warnow T (2018). To include or not to include: the impact of gene filtering on species tree estimation methods. Systematic Biology 67, 285–303.

Mongiardino Koch N (2021). Phylogenomic subsampling and the search for phylogenetically reliable loci. Molecular Biology and Evolution 38, 4025–4038.

Mossel E, Roch S (2008). Incomplete lineage sorting: consistent phylogeny estimation from multiple loci. IEEE/ACM Transactions on Computational Biology and Bioinformatics 7, 166–171.

Nguyen LT, Schmidt HA, Von Haeseler A, Minh BQ (2015). IQ-TREE: a fast and eFective stochastic algorithm for estimating maximum-Likelihood phylogenies. Molecular Biology and Evolution 32, 268–274.

Ogilvie HA, Bouckaert RR, Drummond AJ (2017). StarBEAST2 brings faster species tree inference and accurate estimates of substitution rates. Molecular biology and evolution 34, 2101–2114.

Pamilo P, Nei M (1988). Relationships between gene trees and species trees. Molecular Biology and Evolution 5, 568–583.

Patané JS, Martins Jr J, Setubal JC (2024). A Guide to Phylogenomic Inference. Comparative Genomics: Methods and Protocols, 267–345.

Patel S, Kimball RT, Braun EL (2013). Error in phylogenetic estimation for bushes in the tree of life. Journal of Phylogenetics and Evolutionary Biology 1, 1–10.

Phifer-Rixey M, Harr B, Hey J (2020). Further resolution of the house mouse (Mus musculus) phylogeny by integration over isolation-with-migration histories. BMC evolutionary biology 20, 120.

Price MN, Dehal PS, Arkin AP (2009). FastTree: computing large minimum evolution trees with profiles instead of a distance matrix. Molecular Biology and Evolution 26, 1641–1650.

Price MN, Dehal PS, Arkin AP (2010). FastTree 2–approximately maximum-likelihood trees for large alignments. PLOS ONE 5, e9490.

Rannala B, Yang Z (2003). Bayes estimation of species divergence times and ancestral population sizes using DNA sequences from multiple loci. Genetics 164, 1645–1656.

Rannala B, Yang Z (2017). EPcient Bayesian species tree inference under the multispecies coalescent. Systematic biology 66, 823–842.

Rasmussen MD, Kellis M (2012). Unified modeling of gene duplication, loss, and coalescence using a locus tree. Genome Research 22, 755–765.

Robinson DF, Foulds LR (1981). Comparison of phylogenetic trees. Mathematical Biosciences 53, 131–147.

Sayyari E, Mirarab S (2016). Anchoring quartet-based phylogenetic distances and applications to species tree reconstruction. BMC Genomics 17, 101–113.

Simmons MP, Sloan DB, Gatesy J (2016). The eFects of subsampling gene trees on coalescent methods applied to ancient divergences. Molecular Phylogenetics and Evolution 97, 76–89.

Slatkin M, Pollack JL (2006). The concordance of gene trees and species trees at two linked loci. Genetics 172, 1979–1984.

Song S, Liu L, Edwards SV, Wu S (2012). Resolving confict in eutherian mammal phylogeny using phylogenomics and the multispecies coalescent model. Proceedings of the National Academy of Sciences 109, 14942–14947.

Spielman SJ, Wilke CO (2015). Pyvolve: a fexible Python module for simulating sequences along phylogenies. PloS one 10, e0139047.

Stamatakis A (2014). RAxML version 8: a tool for phylogenetic analysis and post-analysis of large phylogenies. Bioinformatics 30, 1312–1313.

Taillon-Miller P, Bauer-Sardiña I, Saccone NL, Putzel J, Laitinen T, Cao A, Kere J, Pilia G, Rice JP, Kwok PY (2000). Juxtaposed regions of extensive and minimal linkage disequilibrium in human Xq25 and Xq28. Nature genetics 25, 324–328.

Tavaré S (1986). Some probabilistic and statistical problems on the analysis of DNA sequence. Lecture of Mathematics for Life Science 17, 57.

Vachaspati P, Warnow T (2015). ASTRID: accurate species trees from internode distances. BMC Genomics 16, 1–13.

Wang Z, Liu KJ (2016). A performance study of the impact of recombination on species tree analysis. BMC Genomics 17, 165–174.

Wu YC, Rasmussen MD, Bansal MS, Kellis M (2014). Most parsimonious reconciliation in the presence of gene duplication, loss, and deep coalescence using labeled coalescent trees. Genome Research 24, 475–486.

Yan Z, Smith ML, D. P, Hahn MW, Nakhleh L (2022). Species tree inference methods intended to deal with incomplete lineage sorting are robust to the presence of paralogs. Systematic Biology 71, 367–381.

Yin J, Zhang C, Mirarab S (2019). ASTRAL-MP: scaling ASTRAL to very large datasets using randomization and parallelization. Bioinformatics 35, 3961–3969.

Zhang C, Mirarab S (2022a). ASTRAL-Pro 2: ultrafast species tree reconstruction from multi-copy gene family trees. Bioinformatics 38, 4949–4950.

Zhang C, Mirarab S (2022b). Weighting by gene tree uncertainty improves accuracy of quartet-based species trees. Molecular Biology and Evolution 39, msac215.

Zhang C, Rabiee M, Sayyari E, Mirarab S (2018). ASTRAL-III: polynomial time species tree reconstruction from partially resolved gene trees. BMC Bioinformatics 19, 15–30.

Zhang C, Scornavacca C, Molloy EK, Mirarab S (2020). ASTRAL-Pro: quartet-based species-tree inference despite paralogy. Molecular Biology and Evolution 37, 3292–3307.

Zhu T, Flouri T, Yang Z (2022). A simulation study to examine the impact of recombination on phylogenomic inferences under the multispecies coalescent model. Molecular Ecology 31, 2814–2829.

